# Pooled CRISPR screening of high-content cellular phenotypes by ghost cytometry

**DOI:** 10.1101/2023.01.26.525784

**Authors:** Asako Tsubouchi, Yuri An, Yoko Kawamura, Yuichi Yanagihashi, Yuri Murata, Kazuki Teranishi, Soh Ishiguro, Hiroyuki Aburatani, Nozomu Yachie, Sadao Ota

**Author notes:** These authors contributed equally to this work (Asako Tsubouchi, Yuri An, Yoko Kawamura).

## Abstract

Fast enrichment of cells based on morphological information remains a challenge, limiting genome-wide perturbation screening for diverse high-content phenotypes of cells. Here we show that multi-modal ghost cytometry-based cell sorting is applicable to fast pooled CRISPR screening for both fluorescence and label-free high-content phenotypes of millions of cells. By employing the high-content cell sorter in the fluorescence mode, we enabled the genome-wide CRISPR screening of genes that affect NF-κB nuclear translocation. Furthermore, by employing the multi-parametric, label-free mode, we performed the large-scale screening to identify a gene involved in macrophage polarization. Especially the label-free platform can enrich target phenotypes without invasive staining, preserving untouched cells for downstream assays and unlocking the potential to screen for the cellular phenotypes even when suitable markers are lacking.

**One-Sentence Summary:** Machine vision-based cell sorter enabled genome-wide perturbation screens for high-content cell phenotypes even without labeling

## Main Text

CRISPR-based pooled screening has several advantages over conventional array-based approaches for genome-wide perturbation screens: notably increased throughput, reduced cost, and smaller well-to-well batch effect (*1*) (*2*). In the pooled phenotypic screening, cells and intracellular molecules have been labeled with fluorescent dyes, reporters, or immunofluorescent antibodies. Cell phenotyping typically requires the quantification of explicitly defined features where fluorescence-based labeling is advantageous due to its high specificity and sensitivity to the molecules of interest (*3*). For example, representative values such as total fluorescence are measured from temporal signals obtained in fluorescence-activated cell sorting (FACS), or further detailed features such as molecular localization and morphologic parameters are evaluated from optical microscopic images (*4*) (*5*) (*6*) (*7*) (*8*) (*9*). On the other hand, when suitable biomarkers or staining methods are not available and cell phenotypes can only be assessed without labeling, image analysis based on human-recognizable features can become challenging. To solve this issue, machine learning-based analysis of the label-free high-content cell phenotypes is an emerging, promising approach beyond the bias of human recognition (*10*) (*11*).

Here, we present a versatile approach to genome-wide pooled CRISPR screenings for not only fluorescence but also label-free high-content cell phenotypes using a machine vision-based cell sorter (MViCS), which we newly developed based on our ghost cytometry (GC) technologies. The heart of GC is a machine learning-based direct and integrative analysis of cellular morphological information without image production (*11*) (*12*). The high-content information is obtained by flowing cells through a structured light illumination and simultaneously detecting multiple different ghost motion imaging (GMI) waveforms in a temporal domain through different optical paths: fluorescence GMI (flGMI) waveforms excited by a 488 nm laser as well as forward scattering GMI (fsGMI), backscattering GMI (bsGMI), diffractive GMI (dGMI), and bright field GMI (bfGMI) waveforms are generated by a 405 nm laser are measured as analogs to their corresponding microscopic images (Fig. 1A left, see Methods). In the development of a classifier model based on a support vector machine (SVM), we first define the target high-content phenotypes by using the multimodal GMI waveforms with ground truth labels as a training data set. After training, the classifier becomes able to predict the labels directly from the GMI waveforms based on SVM-based scoring. Herein the SVM scores provide users with estimated sorting performance such as a precision-recall (PR) curve as well as an area under the receiver operating characteristic curve (ROC-AUC) score (Fig. 1A right, see Methods).

**Fig. 1.**
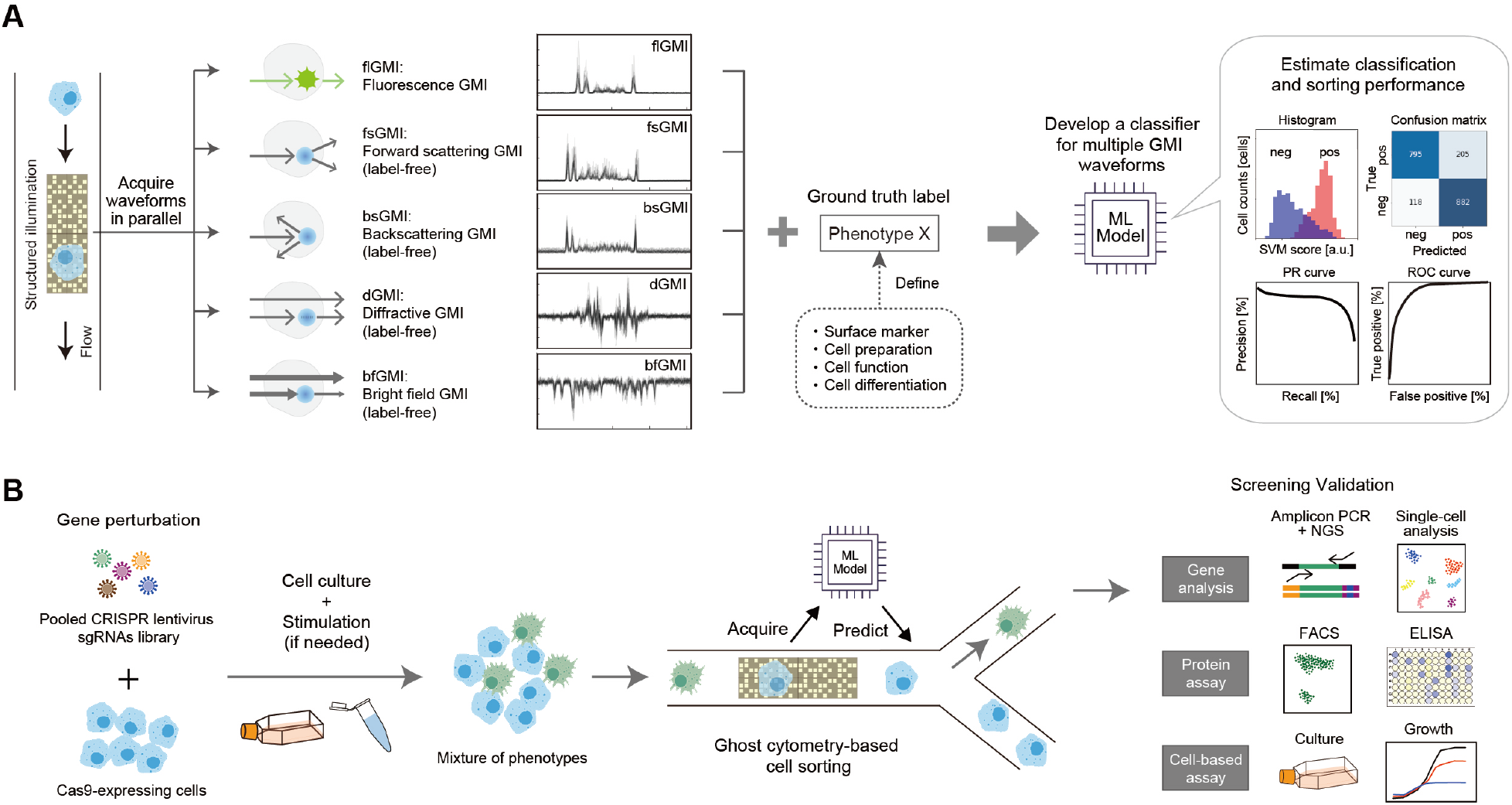
A machine vision-based cell sorter (MViCS) employing multimodal ghost cytometry enabled fast pooled CRISPR screening of high-content fluorescence as well as label-free cell phenotypes. (**A**) Schematic depicting a process of simultaneously acquiring a list of cellular morphological and structural information as different ghost motion imaging (e.g., flGMI, fsGMI, bsGMI, dGMI, and bfGMI) waveforms for each cell and using these data sets for developing machine learning (ML)-based classifier of high-content cellular phenotypes. The GMI waveforms are analogs to microscopic images and used to phenotype cells at subcellular resolutions. (**B**) A workflow of pooled high-content CRISPR screening by the MViCS. A gene-knockout cell library prepared by using CRISPR-Cas9 system is treated with a compound or other reagents to display phenotypes. The MViCS with a pre-trained machine learning model selectively enriches cells that exhibit the target high-content phenotype for downstream analysis. For gene analysis in this work, the sgRNA regions inserted in the isolated genomic DNAs are amplified and sequenced to determine the enriched/depleted genes in response to the treatment. When live cells are used, transcriptomic analysis and cell-based functional assays are widely applicable.

Fig. 1B shows the workflow of CRISPR-based pooled screening using the trained classifier in MViCS. First, cells expressing Cas9 protein are transduced with pooled CRISPR lentiviral libraries for loss-of-function of gene sets and selected for stable viral integration. Next, the pooled knockout cell library is treated with compounds or reagents and displays a variety of phenotypes, where additional assays such as immunostaining can be performed if necessary. We then apply MViCS with a machine learning model trained in advance to the library for selectively enriching cells that exhibit desired high-content phenotypes. Finally, the sorted cells can be subject to various types of biological assays including gene analysis such as genome sequencing, protein assays, and cell-based functional analysis. In the case of a standard CRISPR perturbation screen, after extracting genomic DNAs from the sorted cells, the regions of single-guide RNAs (sgRNAs) are amplified by polymerase chain reaction (PCR), pooled, and then read by commercially available NGS platforms to identify genes that induced the target phenotype. When live cells are sorted, transcriptomic analysis using single-cell RNA sequencing methods as well as cell-based functional assays becomes widely applicable.

We first investigated the capability of MViCS to assess the range of fluorescence high-content cellular phenotypes (Figs. 2). Figs. 2A to D shows example images of an intracellular phenotype, nuclear translocation of NF-κB protein in THP-1 cells, and procedure for developing and evaluating its classifier in MViCS. In the training of a classifier for nuclear translocation, we prepared a cell mixture consisting of LPS-stimulated and unstimulated THP-1 cells stained with the combination of anti-NF-κB primary antibody and Alexa Fluor 488-conjugated secondary antibody; we stained only the unstimulated cells with a ground truth marker dye before mixing them. Note that the total fluorescence intensity for NF-κB staining was similar between the LPS-stimulated and unstimulated cells in the training sample; the two phenotypes were difficult to distinguish using a conventional FACS (Fig. 2B). Instead, we trained the SVM-based classifier for flGMI waveforms representing the nuclear translocation phenotypes using 1,250 cells for each ground truth label. When the trained model is applied to a data set comprising 1,000 waveforms, the histogram of returned scores showed bimodal peaks colored based on ground truth labels, resultantly showing robust and high performance with an AUC score of 0.98 (Figs. 2C, 2D, and fig. S1). Similarly, we tested the capability of MViCS to distinguish the fluorescence distribution in different organelles of adherent HEK293 cells (Figs. 2E to H, and fig. S2) and TFE3 protein subcellular localization in adherent HAP1 parent and *FLCN*-KO cells (*13*) (Figs. 2, I to L, and fig. S3). As a result, the models for each case exhibit high performances with AUC scores of 0.96 and 0.90. The results show that MViCS is able to classify various types of fluorescence high-content phenotypes.

**Fig. 2.**
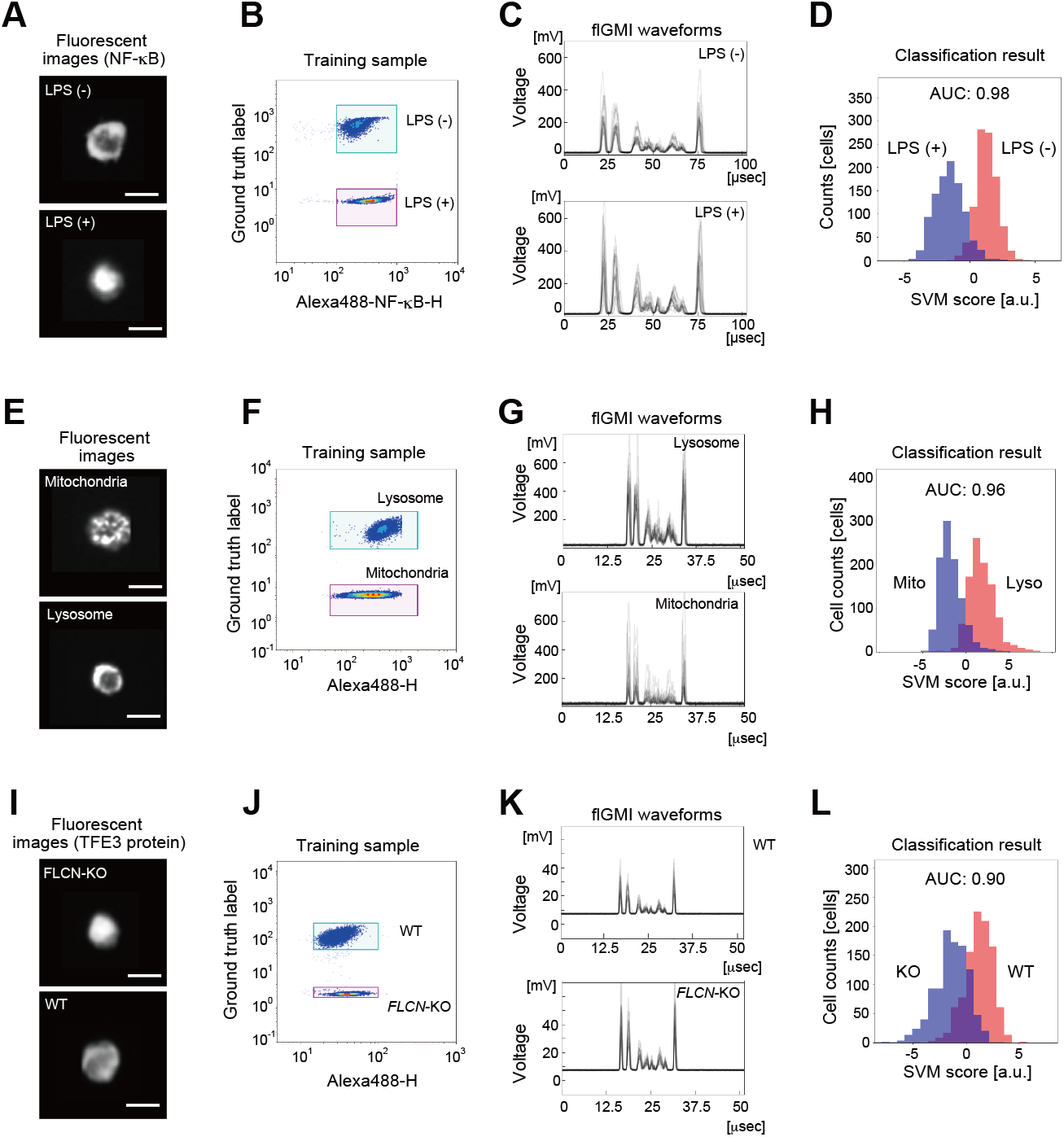
High-content cell phenotypes distinguishable by fluorescent GC. Fluorescence GC classified (**A-D**) nuclear translocation of NF-κB proteins in suspension THP-1 cells, (**E-H**) lysosome (Lamp1) and mitochondria (COX III) in adherent HEK293 cells, and (**I-L**) TFE3 protein subcellular localization in adherent HAP1 parent and *FLCN*-KO cells. (**A, E, I**) Cell images obtained with a commercial imaging flow cytometer are shown left. Scale bars are 10 μm. (**B, F, J**) Conventional FACS scatter plot of fluorescent intensity of labeling to detect cellular phenotyping versus that of a ground truth marker. (**B**) Fluorescent intensity of Alexa488-labeled NF-κB proteins versus that of Fixable Far-Red dye which labeled only LPS unstimulated cells as ground truth. **(F**) Fluorescent intensity of Alexa488-labeled Lamp1 (lysosome) and COX III (mitochondria) proteins versus that of Fixable Far-Red dye which labeled only cells with the Lamp1 proteins stained as ground truth. **(J**) Fluorescent intensity of Alexa488-labeled TFE3 proteins versus that of Fixable Far-Red dye which labeled only parent (WT) cells as ground truth. (**C, G, K**) Example fluorescence GMI (flGMI) waveforms of 20 cells randomly selected for each condition. (**D, H, L**) Classification results of fluorescent cell phenotypes. The performances of ML-based classification are shown as histograms of SVM scores, wherein red and blue colors are assigned by using ground truth labels, and Area Under the ROC curve (AUC) scores.

Herein, we focus on nuclear translocation as the target fluorescence high-content phenotype for the pooled CRISPR screening. NF-κB molecules downstream of the Toll-like receptor 4 (TLR4) pathway are translocated to the nucleus upon the activation by lipopolysaccharide (LPS) stimulation (*14*) (Fig. 3A). As an initial evaluation, we first applied a trained classifier in MViCS for a small-scale pooled cell library in which 60 genes including downstream of the TLR4 pathway were perturbed by using the CRISPR-based system. We prepared 40 sgRNAs targeting 10 genes downstream of the TLR4 signal pathway as positive controls and 20 sgRNAs outside of non-TLR4 signaling pathways as negative controls (table. S1). The cells exhibiting suppression of NF-κB nuclear translocation were sorted and processed for subsequent deep sequencing. The result successfully shows a significant enrichment of cells containing sgRNAs downstream of the TLR4 pathway in the sorted samples (Fig. 3B, figs. S4, and S5). We then conducted the screening at a larger scale using 7,290 sgRNAs targeting 729 kinase genes. We ran 6,000,000 cells through the system and sorted the target cells within 2 hours (S6 and S7). Analysis of the deep sequencing data shows that enriched cells contained sgRNAs targeting *MAP3K7, IRAK4, IKBKB,* and *IKBKG* genes, which are downstream of the TLR4 pathway (Fig. 3C, fig. S8, and S9). Thus, we have demonstrated that the GC-based pooled CRISPR screening is applicable for fluorescent high-content phenotypes at a large scale.

**Fig. 3.**
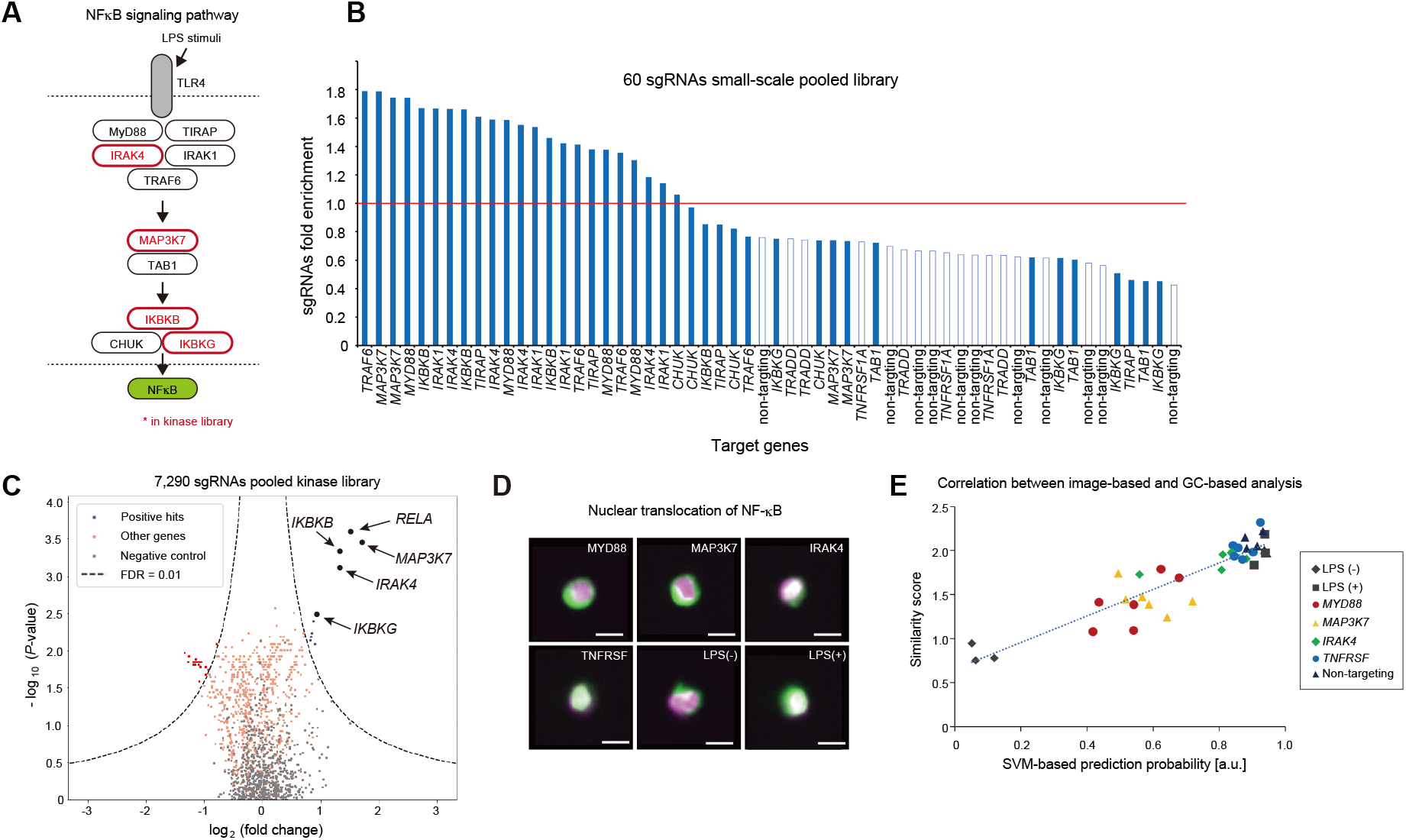
High-throughput pooled CRISPR screening of fluorescent high-content phenotypes. (**A**) Genes downstream TLR4 signaling pathway were targeted to be knocked out by using a small-scale library. **(B)** Enrichment of gRNAs after MViCS-based enrichment based on nuclear translocation of NF-κB. (**C**) Volcano plot visualization of statistical significance (y-axis) and magnitude of the change (x-axis) between before and after the cell sorting, wherein the statistical significance was calculated with Mann-Whitney U test. Dashed lines: cutoff for hit genes (FDR: False discovery rate = 0.01). **(D)** Fluorescent images of NF-κB (green) co-localization with nuclei (magenta) inside *MYD88, MAP3K7, IRAK4,* and TNFRSF CRISPR-knockout cells and those of LPS (-) and LPS (+) cells for control (scale bars are 10 μm). **(E)** A correlation coefficient between SVM-based prediction probabilities in GC and similarity scores obtained by a conventional image flow cytometry was 0.914. *n* = 3 biological replicates.

In addition, we tested whether the degree of cellular phenotype can be quantified using fluorescence GC to the same extent as image analysis using microscopy. Concretely, we compared SVM-based analysis of GMI waveforms and nuclear translocation scores obtained by analyzing images taken with a commercial image flow cytometer for CRISPR knockout cell lines individually targeting *MYD88, MAP3K7, IRAK4,* and *TNFRSF,* respectively. SVM-based prediction probabilities were calculated as values ranging from 0 to 1 using the trained SVM classifier (see Methods), and nuclear translocation scores were obtained as the degree to which two images (i.e., NF-κB and nuclear images) are overlapped using analysis software equipped with the image flow cytometer (*15*) (*16*). The correlation coefficient between these scores was high (R = 0.914) and this result suggests that GC-based phenotypic screening is comparably quantitative as the high-content microscopic image analysis (Figs. 3D, 3E, and fig. S10).

Again, we first investigated the capability of MViCS to assess the range of label-free high-content cellular phenotypes: cell polarization, cell differentiation, and cell exhaustion (Figs. 4). Figs. 4A and 4B show example microscopic images and average scattering properties of label-free morphological phenotypes associated with the polarization of THP-1 cells from M0 to M1 macrophages, respectively. Figs. 4C and 4D show a procedure for developing and evaluating their morphological classifier in MViCS. In the training of a classifier in MViCS, we separately prepared M0 and M1 macrophages and then mixed the two populations as the training sample (*17*) (fig. S11). Herein we used the combination of FSC, BSC, and the label-free fsGMI and bfGMI waveforms to train SVM-based classifiers with 1,000 cells for each ground truth label. When the trained model is applied to a data set comprising 1,000 waveforms, the histogram of returned scores showed bimodal peaks colored based on validation labels, resultantly showing a high performance with an AUC score of 0.89, enabling the classification of the high-content phenotypes of M0 and M1 macrophages in the absence of surface markers. While we note that M0 and M1 polarized macrophages were apparently not easily distinguishable by brightfield microscopy or FACS (Figs. 4A and 4B), this, in turn, supports the MViCS’s ability of robust and accurate classification. Similarly, we tested the capability of MViCS to classify another label-free high-content phenotype of cell differentiation such as THP-1 monocytes and THP-1-derived macrophages (Figs. 4E to G) as well as that of exhausted (LAG3/PD-1 double positive) and non-exhausted (LAG3/PD-1 double negative) human primary T cells (*17*) (*18*) (Figs. 4H to J). As a result, the models for each case exhibit high performances with AUC scores of 0.94 and 0.92 (Fig. 4G, 4J, figs. S12 and S13). The results show that MViCS is able to classify various types of label-free high-content phenotypes.

**Fig. 4.**
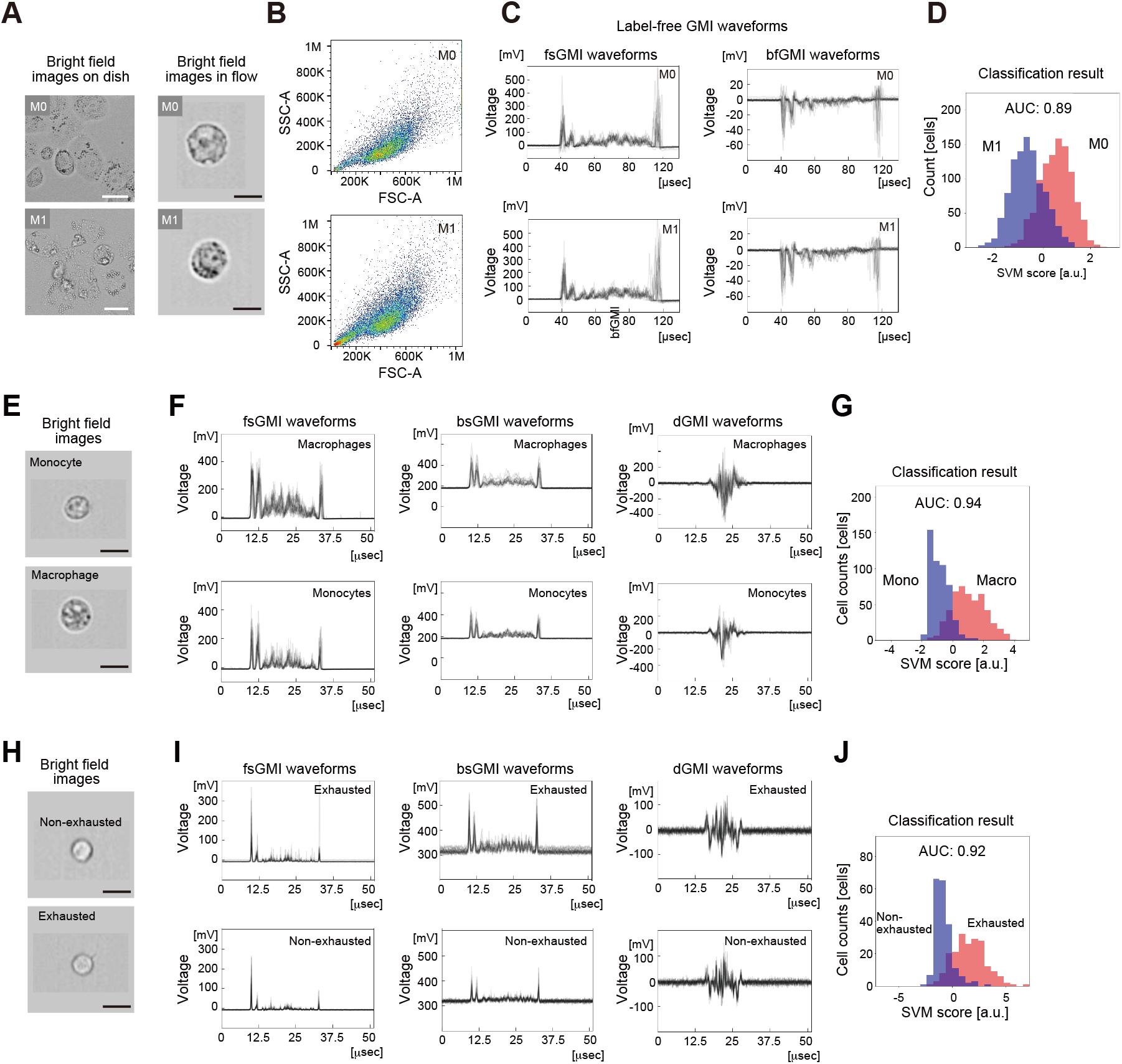
High-content cell phenotypes distinguishable by label-free GC. Label-free GC classified (**A-D**) THP-1-derived M0 and M1 macrophages, (**E-G**) THP-1 monocytes and THP-1-derived macrophages, and (**H-J**) exhausted (LAG3/PD-1 double positive) and non-exhausted (LAG3/PD-1 double negative) human primary T cells. **(A, E, H)** Bright-field cell images on a dish were obtained with a microscope and those in a flow were obtained with a commercial imaging flow cytometer. Scale bars are 30 μm and 10 μm, respectively. (**B**) Conventional FACS scatter plot of FSC and SSC. (**C, F, I**) Example label-free GMI waveforms of 20 cells randomly selected for each condition. (**D, G, J**) Classification results of label-free cell phenotypes. The performances of ML-based classification are shown as histograms of SVM scores, wherein red and blue colors are assigned by using ground truth labels, and Area Under the ROC curve (AUC) scores.

Herein, we focus on the morphological change in the polarization from inactivated macrophage (M0 macrophage) to classically pro-inflammatory macrophage (M1 macrophage) as the target label-free phenotypes and apply the pooled CRISPR screening to identify the genes involved. Macrophages are key cell types in innate immune systems such as tissue repair, inflammation, and cancer, and polarize to different subtypes with various functions including the ability of cytokine secretion and that to respond to injury or pathogenic damage (*19*) (*20*). It is often difficult to define their biological functions with only a few surface markers such that the isolation of live cell populations based on their functions becomes challenging. Thus, assuming that the macrophage polarization correlates with their morphology, we here intend to test if our systems can screen the involved genes based on the change in the label-free high-content cell morphological information without relying on any surface markers. In the training of a classifier in MViCS, we separately prepared M0 and M1 macrophages and then mixed the two populations as the training sample; only M0 macrophages were stained as a ground truth label before mixing with the M1 macrophages. Using the trained classifier and the kinase library, we conducted a large-scale pooled CRISPR screening, where the cells exhibiting suppression of M1 polarization phenotype were sorted and subjected to deep sequencing (Fig. 4A to D, fig. S11, S14 and S15).

By analyzing the enrichment of sgRNA after the sorting, we identified several genes that potentially induced macrophage polarization (Fig. 5A, fig. S16, S17, table S2, and table S3). Notably, the top hit gene, *BRD2*, has been reported as an essential gene for pro-inflammatory cytokine production in macrophages (*17*) (*21*). The results of decreased expression of the M1 marker and secretion of pro-inflammatory cytokines such as IFNγ and TNFα in *BRD2* CRISPR knockout M1 cells supported the hypothesis that the *BRD2* gene is an important modulator of the M1 inducer gene (Figs. 5B, 5C, and fig. S18). Thus, we demonstrated that the large-scale GC-based pooled CRISPR screening is applicable for label-free high-content phenotypes, which can be difficult to distinguish with human-defined features by using standard FACS and possibly even conventional microscopes.

**Fig. 5.**
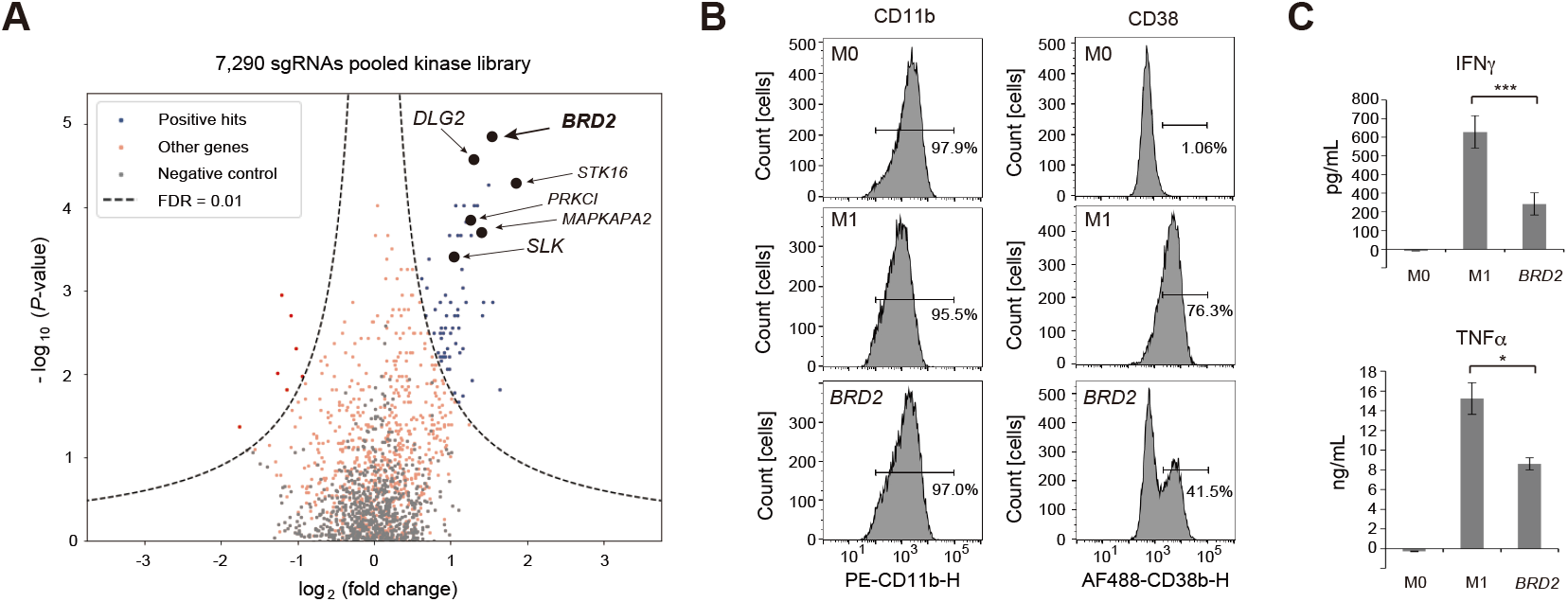
High-throughput pooled CRISPR screening of label-free high-content phenotypes. **(A)** Volcano plot visualization of statistical significance (y-axis) and magnitude of the change (x-axis) between before and after the cell sorting, wherein the statistical significance was calculated with Mann-Whitney U test. Dashed lines: cutoff for hit genes (FDR = 0.01). (B) Expression of macrophage surface markers in control M0, control M1, and *BRD2* CRISPR KO M1 cells, where we used CD11b as a pan-macrophage marker and CD38 as an M1-specific marker. *n* = 3 biological replicates. (C) Cytokine (IFNγ and TNFα) release profiling of the control M0, control M1, and *BRD2* CRISPR KO M1 cells. Supernatants were collected from three independent experiments. Data are presented as mean ± SD; Welch’s t-test: *p < 0.05, ***p < 0.001. *n* = 3 biological replicates.

Machine learning models trained in GC can flexibly target a variety of high-content cell phenotypes, depending on the characteristics of the cell population and the classification objective. In this study, we have trained models with each pair of two cellular phenotypes explicitly defined with a ground truth marker and demonstrated the high performances of these classification models. This approach is extensible to cases where only one phenotype can be defined in the screening process by employing anomaly detection methods such as one-class SVM, which can be readily implemented in the current MViCS. However, there are often cases where no appropriate molecular markers or staining methods are available to define a cell phenotype and the cells of interest show morphological heterogeneity in the pooled population. In such cases, it becomes important to be able to define and sort the desired phenotype by using only the GMI waveforms. To investigate this potential capability of MViCS, we visualized GMI waveforms shown in Figs. 2C and 4C with uniform manifold approximation and projection (UMAP). We performed UMAP after reducing the dimensionality of GMI waveforms with principal component analysis (*22*). (Figs. 6). In Fig. 6A, the UMAP projection of fluorescence GMI waveforms obtained for the mixture of LPS-stimulated and unstimulated cells containing fluorescently labeled NF-κB molecules exhibit clearly distinct two clusters; we confirmed that the two are LPS-stimulated and unstimulated ones, respectively. The distinct representation of LPS-stimulated and unstimulated populations indicate that we can potentially train classification models based on the populations defined in the UMAP space as we do for molecular marker-based populations. In addition, Fig. 6B shows that cells represented as distinct in the UMAP space have different SVM scores, which were quantified in Fig. 2D. This result indicates that morphological differences identified in supervised learning algorithms can also be captured by unsupervised learning algorithms including UMAP and other dimensionality reduction methods. In Fig. 6C, same as the fluorescence case, we projected the label-free GMI waveforms obtained for the mixture of THP-1-derived M0 and M1 macrophage cells in UMAP with colors assigned based on the ground truth labels. As a result, Although M0 and M1 macrophages were not represented as distinct clusters in the UMAP projection, Fig. 6D shows that their distributions are still consistent with the SVM score obtained from the classification model in Fig. 4D. These analysis results strongly support that the GMI waveforms alone will allow us to define the target high-content cellular phenotypes if the morphological difference is discernible in unsupervised manners.

**Fig. 6.**
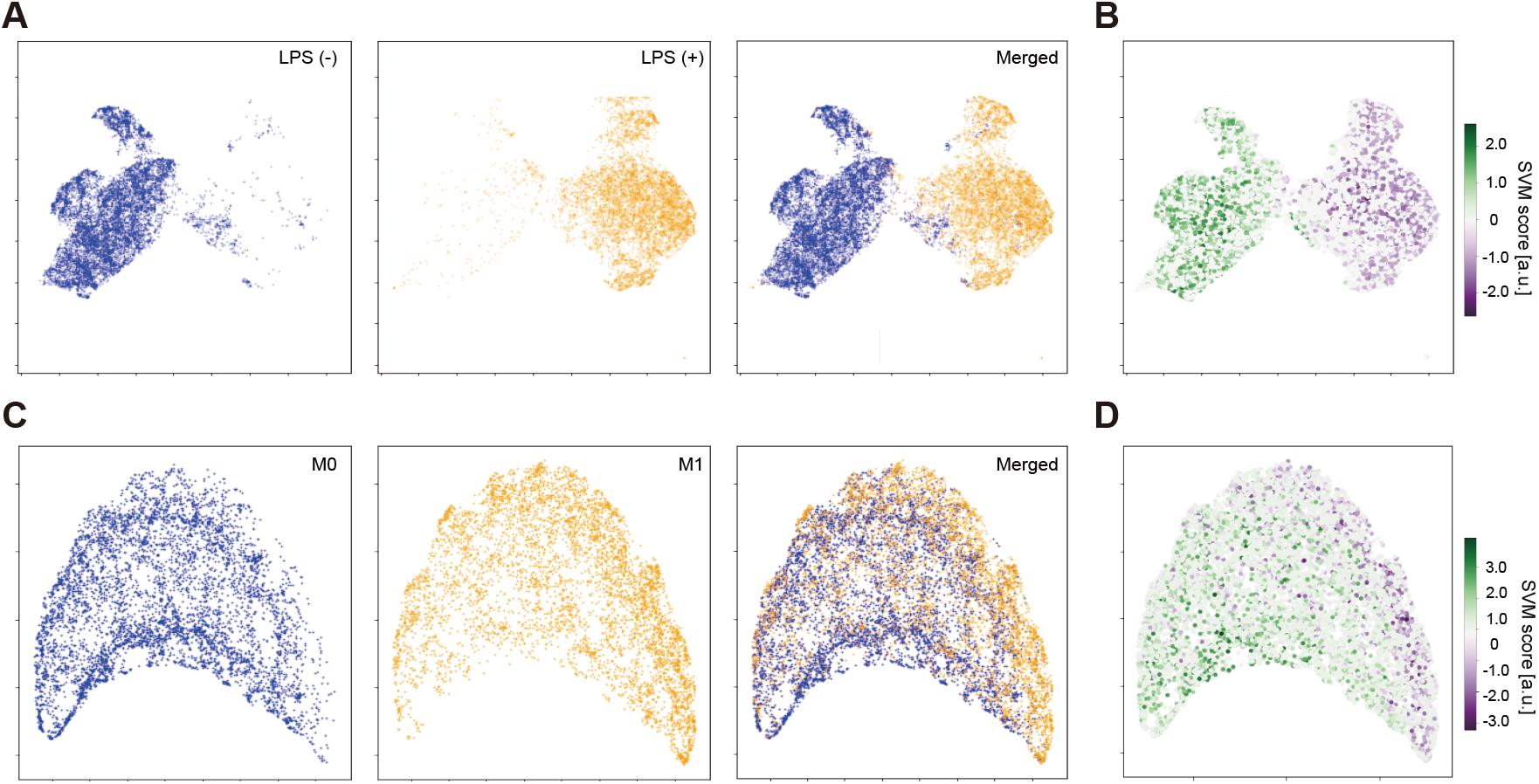
UMAP projection of GMI waveforms. (**A**) UMAP projection of fluorescence GMI waveforms obtained for LPS stimulated (orange) and unstimulated THP-1 cells (blue). The same UMAP plot shown in **A** is colored according to SVM scores obtained in the classification of NF-kB fluorescence GMI waveforms for the LPS-treated cells (stimulated vs unstimulated). **(C)** UMAP projection of label-free GMI waveforms obtained for THP-1-derived M0 (blue) and M1 (orange) macrophage cells. **(D)** The same UMAP plot shown in **C** is colored according to SVM scores obtained in the classification of label-free multimodal GMI waveforms for the cells at different polarization states (M0 vs M1).

A technical advantage of MViCS-based CRISPR screening over conventional FACS-based screening is that it integratively analyzes more detailed information about cells for a more accurate cell selection. In fluorescence-based cell phenotyping, this method is shown to be powerful when there is no significant difference in total fluorescence intensity despite different cell phenotypes, such as subcellular localization of proteins or changes in different organelles (Figs. 2). The advantage of utilizing high-dimensional GMI waveforms is also true for the case of label-free cellular phenotyping (Figs. 4). Indeed, only with conventional FSC and SSC (Fig. 4B), the classification of macrophage polarization states becomes difficult even if the SVM method is used (fig. S11D). The analysis in fig. S11D also shows that the combinatorial use of the average characteristics of FSC and SSC with the detailed GMI waveforms constructively improves the classification accuracy. Similarly, we can also foresee that the combinatorial use of surface markers-based cell definition with the label-free high-content cellular phenotyping in MViCS is beneficial to look at finer cell differences between cell subtypes of immune cells and cell states such as activation, exhaustion, and differentiation.

MViCS’s unique capability of resolving both the fluorescence and label-free high-content cellular phenotypes holds potential for the discovery of various new targets including gene perturbations as well as compounds. The fluorescence mode is effective, especially when molecules or subcellular features of interest are predetermined, and enables the screening based on the change in the spatial distribution of the targeted molecules or features. Examples include aggregation or degradation of proteins as well as decomposition of intracellular organelles as indicators of mechanism of action (*23*) (*24*) (*25*). The label-free GMI waveforms uniquely capture the detailed morphological change as a phenotypic response of each entire cell. We further prospect that, in future implementations, the simultaneous use of the fluorescence and label-free GMI waveforms will enable cell-based screening based on the combinatorial analysis of the changes in fluorescently labeled target proteins and the detailed holistic high-content cell phenotypes (*26*) (*27*) (*28*).

In summary, we report the development of high-content and large-scale pooled CRISPR screening by fast and selective sorting of cells based on machine vision on their high-content phenotypes. We anticipate that MViCS will be widely used for the screening of further important cellular phenotypes by combining various available machine learning methods and existing biomarkers with its high-content analysis capability. When combined with single-cell sequencing techniques (*29*) (*30*) (*31*) (*32*), this method is compatible with the pooled screening of various DNA-tagged perturbations including antibodies (*33*), compounds (*34*), shRNAs (*35*) and peptides (*36*). Importantly, not limited to the fluorescence-based cellular phenotypes, our machine learningbased label-free high-content cell analysis enabled the enrichment of target phenotypes without invasive staining, preserving ‘untouched’ cells for downstream functional assays and thus finding a wide range of biological applications.

## Acknowledgments

We thank all members of ThinkCyte Inc. for experimental support and discussions, in particular Keisuke Toda, Yasuhiro Kajiwara, Hikari Morita, and Keiji Nakagawa for the development of the instrument, Hiroaki Adachi for supporting the data analysis, Hiroaki Suita and Kaoru Komoriya for supporting biological experiments, Keisuke Wagatsuma and Hirofumi Nakayama for scientific discussions. Greg Schneider, Andy Wu, and Willem Westra for reviewing the manuscript. We also thank Kaori Shiina from the University of Tokyo for supporting the analysis of sequencing data.

## Funding

This work was supported by the Product Commercialization Alliance (PCA) program funded by the New Energy and Industrial Technology Development Organization (NEDO).

## Author contributions

A.T. and S.O. conceived the study, designed the experiments, interpreted the results, and wrote the manuscript. S.O. supervised the study. A.T. developed experimental tools and performed most experiments; construction of CRISPR library, flow cytometry analysis, cell culture, cell staining, data analysis of ghost cytometry, and NGS library preparation. Y.A. performed experiments and data analysis of ghost cytometry and NGS. Y.K. designed and fabricated microfluidic sorting chips, integrated the GC-based cell sorter, and performed all the GC-based cell sorting experiments. Y.M. performed all the GC-based cell sorting experiments. Y.Y. performed computational analysis of NGS data analysis. T.K. performed ghost cytometry analysis. S.I. H.A. and Y.N. provided conceptual input on pooled CRISPR screening, CRISPR library design, library cloning strategy, and technical input. All authors contributed to the revision of the manuscript.

## Competing interests

S.O. is the founder and shareholder of ThinkCyte Inc., a company engaged in the development of ghost cytometry (GC)-based cell sorting. A.T., Y.A., Y.K., Y.M., Y.Y., and T.K. have shares of stock options of ThinkCyte Inc. S.O., A.T., Y.A., Y.K., Y.M., Y.Y., and T.K. have filed patent applications related to this paper.

## Data and materials availability

All data are available in the main text or the supplementary materials. Data analysis was performed with customized code written in Python which will be available on Zenodo.

## Supplementary Materials

Materials and Methods

Figs. S1 to S18

Tables S1 to S4

