## Supplementary Materials for "Pooled CRISPR screening of high-content cellular phenotypes by ghost cytometry"

### Materials and Methods:

#### Cell culture

HEK293T cells were purchased from Applied Biological Materials (abm) and maintained in Dulbecco's Modified Eagle's Medium (DMEM) (FUJIFILM Wako) supplemented with 10 % fetal bovine serum (FBS) (HyClone) and 1 % penicillin-streptomycin solution (FUJIFILM Wako) at 37 °C with 5 % $CO_2$. Cells were routinely tested for mycoplasma contamination by nested PCR using culture medium as a template.

HAP1 parent and *FLCN*-KO cells were purchased from Horizon and maintained in Iscove's Modified Dulbecco's Medium (IMDM) (FUJIFILM Wako) supplemented with 10 % fetal bovine serum (FBS) (HyClone) and 1% penicillin-streptomycin solution (FUJIFILM Wako) at 37 °C with 5 % $CO_2$. Cells were routinely tested for mycoplasma contamination by nested PCR using a culture medium as a template.

THP-1 human monocytic leukemia cells were purchased from ATCC and maintained in RPMI-1640 (FUJIFILM Wako) supplemented with 10 % fetal bovine serum (FBS) (HyClone), 1% penicillin-streptomycin (FUJIFILM Wako), and 1 % StemSure 2-mercaptoethanol solution (FUJIFILM Wako) at 37 °C with 5 % $CO_2$. Cells were routinely tested for mycoplasma contamination by nested PCR using culture medium as a template. For activation of TLR4 signal, THP-1 cells were stimulated with lipopolysaccharide (LPS) (Sigma-Aldrich) (300 ng/mL) for 1 hour at 37 °C with 5 % $CO_2$. For macrophage differentiation and polarization from THP-1 cells, THP-1 monocytes were differentiated into resting macrophages (M0) using 100 nM phorbol 12-myristate 13-acetate (PMA) (Sigma-Aldrich) for 72 hours followed by 24 hours in PMA-free medium. For M1 polarization, M0 macrophages were further cultured in M1-polarization medium containing 100 ng/mL LPS (Sigma-Aldrich) and 20 ng/mL IFNγ (R&D systems) for 24 hours starting on the third day of PMA treatment (*17*).

For primary T cells, Pan-T cells were purchased from Precision for medicine and culture in X-VIVO 15 (Lonza) supplemented 10 % fetal bovine serum (FBS) (HyClone), 1 % Glutamax supplement (Thermo Fisher), 1 % penicillin-streptomycin (FUJIFILM Wako), and 1 % StemSure 2-mercaptoethanol solution (FUJIFILM Wako) at 37 °C with 5 % $CO_2$. To mimic a transient stimulation, cells were cultured in the presence of CD3/CD28 Dynabeads (Thermo Fisher) and 25 U/mL of rhIL-2 (Pepro Tech) for 3 days and then rested in the presence of only rhIL-2 for the remaining 11 days of culture (*18*).

#### Construction of small-scale CRISPR sgRNA expression plasmids

gRNA spacer inserts were prepared by a single pot reaction to phosphorylate and anneal ssDNA pairs. To prepare each spacer fragment (table S1), a T4 polynucleotide kinase reaction sample was prepared with two ssDNAs in accordance with the manufacturer's protocol (Takara) and placed in a thermal cycler with the following conditions: 37 °C for 30 min; 95 °C for 5 min; 70 cycles of 12 sec starting with 95 °C and -1 °C per cycle, and then maintained at 25 °C. The annealed spacer inserts were then ligated into a pLentiGuide-Puro (Addgene) by Golden Gate Assembly using BsmBI (NEB) and T4 DNA ligase (NEB). The assembly was performed under the following thermal cycler conditions: 15 cycles of 37 °C for 5 min and 20 °C for 5 min, 55 °C for 30 min, and then maintained at 4 °C (*37*). pLenti-Cas9-Blast were purchased from Addgene. The plasmid sequences were confirmed by Sanger sequencing.

**Amplicon sequencing**
Genomic DNA in fixed cells was extracted using QIAamp DNA FFPE Tissue Kit (QIAGEN) or hotshot method (*37*) From each sample, the target regions were amplified using the extracted genome DNA as the PCR template with its corresponding first HTS primer pair (table S4). The PCR was performed according to the previous protocol (*37*). The PCR product was extracted using FastGene Gel/PCR Extraction Kit (Nippon Genetics) and reamplified using custom Illumina index primers (table S4). Each indexed library was electrophoresed in a 2 % agarose gel and the expected band was purified using FastGene Gel/PCR Extraction Kit (Nippon Genetics). The sequencing libraries were quantified by qPCR using KAPA Library Quantification Kit Illumina (KAPA Biosystems) for multiplexing. The multiplexed libraries were quantified by the same qPCR protocol and sequenced with 30 - 40% PhiX control using Illumina HiSeq2500 (TruSeq rapid SBS kit; 2 x 151 bp paired-end) or MiSeq (MiSeq v2 kit; 2 x 301 bp paired-end).

**Lentivirus production**
HEK293T cells were seeded into a 10-cm plate at a density of 100,000 cells/cm$^2$. After 20 hours, cells were transfected with pMD2.G (0.75 µg) (Addgene), psPAX2 (2.25 µg) (Addgene), and a lentiviral transfer plasmid (3.6 µg) using polyethylenimine (PEI) MAX (Cosmobio) (*37*). Media was exchanged after 24 hours. Viral supernatant was harvested 48 hours after transfection and filtered through 0.45 µm cellulose acetate filters. CellTiter-Glo (Promega) was used for the measurement of virus titer according to the manufacturer's protocol.

**Lentiviral transduction**
THP-1 cells expressing Cas9 protein were transduced in bulk with the library at low multiplicity of infection (MOI) (0.15 viral particles per cell) by adding viral supernatant supplemented with polybrene (8 µg/mL) and centrifuging at 510 g for 1 hour at 37 °C. After 24 hours post-infection, cells were passaged into media containing selection antibiotic at the following concentrations: 0.5 ug/mL puromycin (Sigma-Aldrich) and 10 ug/mL blasticidin (Sigma-Aldrich).

**Immunostaining**
HEK293T cells were washed with PBS once and treated with 0.25 w/v% Trypsin EDTA solution (FUJIFILM Wako) for 5 min at 37 °C for detaching from the culture dish. Cells were fixed with 4 % formaldehyde in PBS (Thermo Fisher) for 20 min at room temperature. For visualizing mitochondria, cells in PBS were incubated at 95 °C for 5min, blocked with 5 % FSB with 0.3 % Triton X-100 in PBS solution for 30 min at room temperature, and incubated with 1/200 diluted anti-UQCRC1 antibody for Complex III-core1 (Invitrogen, #459140) in the blocking buffer for overnight at 4 °C. After washing with wash buffer (0.5 % FBS in PBS), cells were incubated with Alexa Fluor 488 conjugated secondary antibody (Thermo Fisher, #A32723) for 1 hour at room temperature. For visualizing lysosome, cells were permeabilized with 0.2 % Triton X-100 in PBS for 3 min at room temperature after fixation. After washing with 0.5 % BSA in PBS once, cells were blocked with 5 % BSA in PBS for 1 hour at room temperature and incubated with 1/200 diluted anti-Lamp1 antibody (CST, #9091) in 1 % BSA in PBS overnight at 4 °C. After washing with 0.5 % BSA in PBS, cells were incubated with Alexa Fluor 488 conjugated secondary antibody (Thermo Fisher, #A11008) for 1 hour at room temperature.

HAP1 parent and *FLCN*-KO cells were fixed with 4 % formaldehyde in PBS (Thermo Fisher, #R37814) for 20 min at room temperature. After washing with PBS once, cells were incubated with ice-cold methanol for 5 min at -20 °C. After washing with 0.5 % BSA in PBS, cells were blocked with 5 % BSA in PBS for 1 hour at room temperature and incubated with anti-TFE3 antibody (Sigma Aldrich, #023881) in 1 % BSA in PBS overnight at 4 °C. After washing with 0.5 % BSA in PBS, cells were incubated with Alexa Fluor 488 conjugated secondary antibody (Thermo Fisher, #A11008) for 1 hour at room temperature.

THP-1 cells were fixed with 4 % formaldehyde in PBS (Thermo Fisher, #R37814) for 20 min at room temperature. After washing with PBS once, cells were incubated with ice-cold methanol for 5 min at -20 °C. After washing with 0.5 % BSA in PBS, cells were blocked with 5 % BSA in PBS for 1 hour at room temperature and incubated with anti-NF-κB p65 antibody (CST, #8242) in 1 % BSA in PBS overnight at 4 °C. After washing with 0.5 % BSA in PBS, cells were incubated with Alexa Fluor 488 conjugated secondary antibody (Thermo Fisher, #A11008) for 1 hour at room temperature.

**Flow cytometry**

The following monoclonal antibodies were used for flow cytometry and ghost cytometry: PE anti-human LAG3 (11C3C65), APC anti-human PD-1 (EH12.2H7), PE anti-human CD11b (ICRF4), and Alexa Fluor488 anti-human CD38 (HIT2) along with the corresponding isotype controls (all from BioLegend). Cell viability was assessed by 7-AAD (BD Biosciences) or Zombie NIR staining (BioLegend). Hoechst 33342 (Thermo Fisher, #H3570) was used for nuclear staining. Cells were suspended in stain buffer (2% FSB with 1 mM EDTA in PBS) with human FcR blocking reagent (Miltenyi Biotec, #130-059-901) and incubated with monoclonal antibodies for 30 min at 4 °C. Flow cytometry data was acquired using Attune Flow Cytometer (Thermo Fisher) or JSAN (Bay bioscience). Fluorescence was compared to the corresponding isotype-stained controls. All data were analyzed using the FlowJo software (Tree Star Inc.). Cell images in flow were taken by FlowSight system, and black or gray areas were added to their surroundings to present uniformly sized pictures for better visualization in figures. The similarity scores NF-κB and Hoechst were obtained by IDEAS 6.2 (Luminex).

**ELISA assay**

THP-1 cells were polarized as described above. The supernatant was collected from three independent experiments, pooled, and subjected to the Human Pro-inflammatory Cytokine Multiplex ELISA Kit (Arigo Biolaboratories Corp) according to the manufacturer's protocol. Read the OD with a microplate reader (Speark, Tecan) at 450 nm and calculate the average absorbance values for each set of standards and samples. The experiments were performed as three independent replicates.

**Experimental conditions for fluorescence and label-free GC**

The detail of the electronics setting was described previously (*11*) (*12*). Briefly, all photomultiplier tubes (PMTs) were purchased from Hamamatsu Photonics Inc. PMTs of 10 MHz with built-in amplifiers were used for detecting the flGMI, fsGMI, bsGMI, and dGMI signals, while PMTs of 200 kHz, 1 MHz, or 10 MHz were used to detect fluorescence signals and BSC signals. FSC signals were obtained using either a photodetector or a PMT of 200 kHz. Multi-pixel photon counter from Hamamatsu photonics Inc. was used to detect bfGMI. The direct current of flGMI, fsGMI, bsGMI, bfGMI, dGMI, and FSC signals were cut with an electronic high-pass filter. The PMT signals were recorded with electronic filters using a digitizer or an FPGA development board with a homemade analog/digital converter. The digitizer and/or FPGA continually collected a fixed length of signal segments from each color channel at the same time, with a fixed trigger condition applied to the FSC signals.

The cells were flowed through either a quartz flow cell (Hamamatsu) or a polydimethylsiloxane (PDMS)-based microfluidic device using a customized pressure pump and/or a syringe pump (KD Scientific). The quartz flow cell had a channel cross-section dimension of 150 x 150 μm$^2$ at the measurement position, wherein the sheath fluid (IsoFlow, Beckman Coulter) was driven at a pressure of 305 kPa for the cell analysis. The sample fluid was driven at a flow rate between 10 and 40 μl/min. For cell sorting, custom handmade sorting chips were used. The PDMS device had a channel with a cross-section dimension of 32 x 80 μm2 at the measurement position, wherein the sheath flow was driven at a pressure of about 180 to 270 kPa and the sample fluid was driven at a flow rate of 20 μm/min in the cell sorting. For sorting action, a piezoelectric (PZT) actuator implemented on the sorting chip was driven by an input voltage and displaced a fluid containing target cells toward a collection channel.

The detail of the machine learning method was described previously (*11*) (*12*). For the binary classification of cells, a support vector machine (SVM) algorithm with radial basis function (rbf) kernel included in the scikit-learn library was used. All training and validations of the models were performed using equal amounts of samples for each class label. The number of cells used for training and testing SVM is described in the subsequent sections. After training, probability or decision function was computed using the predict_proba method or the decision_function method from the scikit-learn library, and these were used as SVM scores. Distributions of these scores for each sample were visualized as histograms using the matplotlib library. We evaluated trained machine learning models using accuracy, receiver operating characteristic curve (ROC curve), and area under the ROC curve (AUC or ROC-AUC). ROC curve was drawn using the matplotlib library after calculation of the true positive rate (tpr) and false positive rate (fpr) with the scikit-learn library. Precision-recall pairs for different thresholds of SVM score were computed with the scikit-learn library and the precision-recall curve (PR curve) was drawn using the matplotlib library. The hyperparameters (regularization parameter and kernel parameter in SVM) were optimized by 3-fold cross-validation of the AUC score.

**Classification of lysosome and mitochondria**

HEK293T cells were immunostained with anti-Lamp1 antibody and anti-UQCRC1 antibody described above. As a ground truth label, anti-Lamp1atnibody stained cells were incubated with LIVE/DEAD fixable far red (FFR) dead cell staining kit (Thermo Fisher). For the classification of lysosomal and mitochondrial organelle patterns of fluorescence distribution, each of the cells was mixed at an equal concentration. The mix suspension was allowed to flow through the fluorescence GC system. The cells were gated using FSC/SSC scatter plot to remove doublets and debris from the training sample. flGMI waveforms were used as a classification modality. Within the data, 2,500 and 1,000 cells selected randomly (without overlap) with an equal number of anti-Lamp1 and anti-UQCRC1 stained cells were used as training and testing data, respectively.

**Classification of TFE3 subcellular localization**

HAP1 parent and *FLCN*-KO cells were immunostained with anti-TFE3 antibody described above (*13*). As a ground truth label, HAP1 parent cells were incubated with LIVE/DEAD fixable far red (FFR) dead cell staining kit (Thermo Fisher). For the classification of subcellular localization of TFE3 protein in parent and *FLCN*-KO cells, each of the cells was mixed at an equal concentration. The mix suspension was allowed to flow through the fluorescence GC system. The cells were gated using FSC/SSC scatter plot to remove doublets and debris from the training sample. flGMI waveforms were used as a classification modality. Within the data, 3,000 and 1,000 cells selected randomly (without overlap) with an equal number of parent and *FLCN*-KO cells were used as training and testing data, respectively.

**Classification of THP-1-derived monocytes and macrophages**

THP-1-derived monocyte and macrophages were stained with cell surface markers and the viability dye described above. For the classification of exhausted and non-exhausted cells, the cells were gated using FSC/SSC scatter plot to remove doublets and debris. After gating live cells, CD11b positive cells were identified as the macrophage population and CD11b negative cells were identified as the monocytes population. fsGMI, bsGMI, and dGMI waveforms were used as classification modalities. Within the data, 1,000 and 500 cells selected randomly (without overlap) with an equal number of monocytes and macrophages were used as training and testing data, respectively.

**Classification of exhausted and non-exhausted cells**

Transient stimulated T cells were stained with cell surface markers and viability dye described above. For the classification of exhausted and non-exhausted cells, the cells were gated using FSC/SSC scatter plot to remove doublets and debris. After gating live cells, LAG3/PD-1 double-positive cells were selected as exhausted cells and LAG3/PD-1 double-negative cells were selected as non-exhausted T cell populations. fsGMI, bsGMI, and dGMI waveforms were used as classification modalities. Within the data, 400 and 400 cells selected randomly (without overlap) with an equal number of exhausted and non-exhausted cells were used as training and testing data, respectively.

**Classification and sorting of CRISPR library by fluorescence GC**

THP-1 cells were immunostained with anti-NF-κB antibody described above. As a ground truth label, a population of LPS unstimulated cells was stained with LIVE/DEAD fixable far red (FFR) dead cell staining kit (Thermo Fisher). For the classification of LPS stimulated and unstimulated cell populations, each of the cells was mixed at an equal concentration. The mix suspension was allowed to flow through the fluorescence GC system. The cells were gated using FSC/SSC scatter plot to remove doublets and debris from training data sets. flGMI waveforms were used as a classification modality. Within the data, 1,250 and 1,000 cells (without overlap) with an equal number of LPS stimulated and unstimulated cells were used as training and testing data, respectively.

The pooled CRISPR library cells were stimulated with LPS and stained with anti-NF-κB antibody described above. When a classifier was trained and implemented on an FPGA, it judged the cells in real-time using GMI signals and then enabled subsequent sorting of the cells according to the judgment. The actual time lapsed during the judgment for a single cell with the FPGA was 6.0 μs. For small-scale library sorting, 100,000 cells were applied to the sorter and 70,000 cells were predicted as positive and sorted by the trained model. The experiments were performed as two independent replicates. The cells expressing pooled 40 positive sgRNAs out of 60 total sgRNAs were labeled with a ground truth marker for validation of the sorting purity. To evaluate the actual sorting purity, we counted the number of positive cells in the sorted sample by using a conventional FACS and confirmed that 90.8 % cells in the sorted sample were labeled with the ground truth marker (fig. S4B). The result obtained by the conventional FACS was close to the predicted sorting purity of 92.6 % and recovery of 96.3 % (fig. S4A). The reproducibility of making a trained model was evaluated (the average of the AUC score was 0.98 ± 0.02, $n = 7$) and high Z'-factor value from the SVM score (0.83, $n = 7$) was obtained (*38*) (fig. S4C). For sorting of kinase library, 6,000,000 cells (coverage 500~) were applied to the sorter and 600,000 cells were predicted as positive and sorted by the trained model within 2 hours. We set the threshold for SVM scores more than 0 for cell sorting and the percentage of cells predicted as positive was around 7.6 % (fig. S6F). The experiments were performed as four independent replicates.

**Classification and sorting of CRISPR kinase library by label-free GC**

THP-1-derived M0 and M1 macrophages were prepared individually as described above. After washing PBS once, cells were incubated with 0.25 w/v% Trypsin EDTA solution (FUJIFILM Wako) for 5 min at 37 °C and scraped gently using Cell Lifter (Corning) for detaching from the culture dish. As a ground truth label, M0 macrophages were stained with CellTracker Green CMFDA Dye (Thermo Fisher). For the classification of M0 and M1 macrophages, each of the cells was mixed at an equal concentration. The mix suspension was allowed to flow through the fluorescence GC system. The cells were gated using FSC/SSC scatter plot to remove doublets and debris from a training data set. fsGMI and bfGMI waveforms were used as classification modalities. Within the data, 1,000 and 1,000 cells selected randomly (without overlap) with an equal number of M0 and M1 macrophages were used as training and testing data set, respectively. The reproducibility of making a trained model was evaluated (the average of the AUC score was 0.85 ± 0.06, $n = 7$) and a high Z'-factor value from the SVM score (0.75, $n = 7$) was obtained (*38*) (fig. S14C). For validation of differentiation and polarization, the M0 and M1 macrophages for the training sample were stained with known M0 and M1 CD markers individually (*17*) (figs. S14D to G).

Pooled CRISPR kinase library cells were differentiated and incubated with an M1-polarization medium. For sorting of the kinase library, 6,000,000 (coverage 500~) cells were applied to the sorter, and 150,000 cells predicted as positive were sorted within 2 hours by using a classifier trained and implemented on an FPGA. To increase the purity, we set the threshold of SVM scores of more than 1 for cell sorting. If the percentage of target cells in the pre-sorting sample was 3 %, the predicted purity was 37.5 % and the predicted recovery was 16.7 % in the sorted sample (figs. S14A and S14B). The experiments were performed as three independent replicates.

**Bioinformatic analysis of CRISPR data**

Statistical analysis of sgRNA enrichment in CRISPR screening. Adapter trimming and demultipexing were performed using Ultraplex (*39*). According to the published bioinformatics pipeline, the large-scale screening data were analyzed (*40*) (*41*) (kampmannlab.ucsf.edu/mageck-inc). Briefly, first, raw sequencing reads from next-generation sequencing were cropped and aligned to the reference using Bowtie2 to determine sgRNA counts in each sample. Next, counts files of two samples subject to comparison were input into MAGeCK and $\log_2$ fold changes (LFCs) and *p*-values were calculated for each sgRNA. Gene level knockout phenotypes scores were determined by averaging LFCs of the top 3 sgRNAs targeting this gene with the most significant *p*-values. The statistical significance for each gene was determined by comparing the set of *p*-values for sgRNAs targeting it with the set of *p*-values for non-targeting control sgRNAs using the Mann-Whitney U test. Based on the distribution of all the products, a cutoff value was chosen to make sure the false discovery rate (FDR) is less than 0.01.

Fig. S1

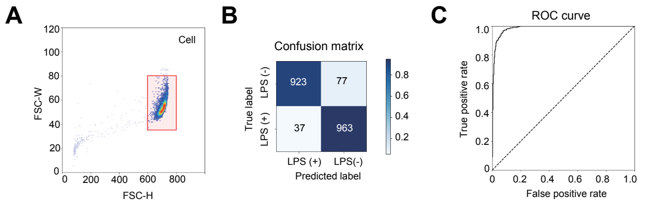


**Fig. S1.**

The development and performance of a machine learning-based model which classifies the high-content phenotypes of fluorescently labeled NF-κB in suspension THP-1 cells based on fluorescence GMI waveforms. **(A)** Gating was first performed by the cell gate. **(B)** Confusion matrix of the classification result. **(C)** ROC curve.

Fig. S2

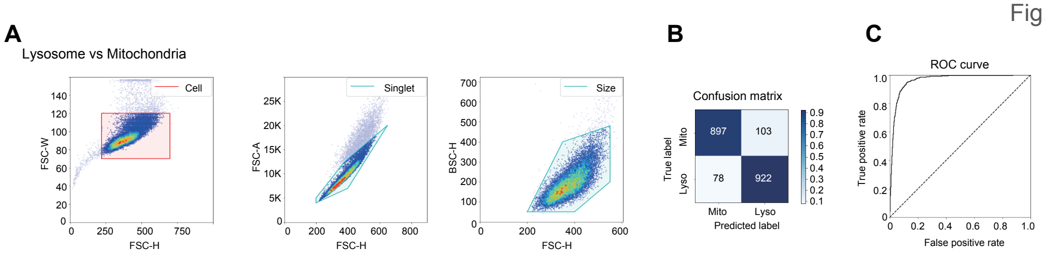


**Fig. S2.**

The development and performance of a machine learning-based model which classifies the high-content phenotypes of fluorescently labeled lysosome (Lamp1) and mitochondria (COX III) in adherent HEK293 cells based on fluorescence GMI waveforms. **(A),** Gating scheme; cell gate, singlet gate, and size gate. **(B)** Confusion matrix of the classification result. **(C)** ROC curve.

Fig.S3

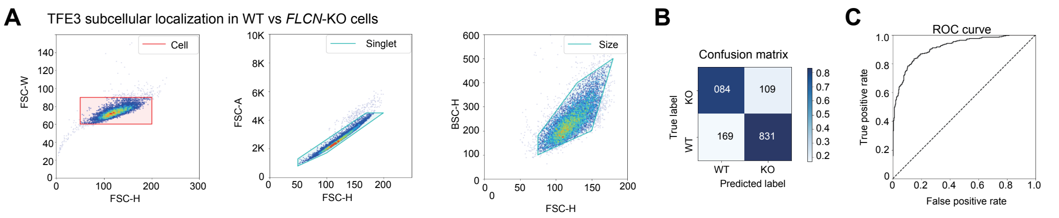


**Fig. S3.**

The development and performance of a machine learning-based model which classifies the high-content phenotypes of fluorescently labeled TFE3 protein subcellular localization in adherent HAP1 parent and *FLCN*-KO cells based on fluorescence GMI waveforms. **(A)** Gating scheme; cell gate, singlet gate, and size gate. **(B)** Confusion matrix of the classification result. **(C)** ROC curve.

Fig.S4

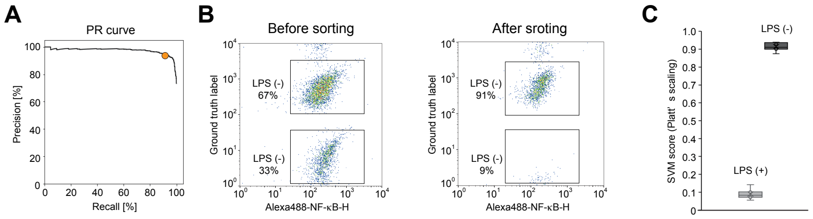


**Fig. S4.**

Data obtained in the pooled CRISPR screening of a small-scale (60 sgRNAs) library by using MViCS in the fluorescence mode (N1: a sample used in Fig. 2A-2D, Fig. 3B). **(A)** PR curve. A Yellow circle indicates the threshold of 0 < SVM score. **(B)** Validation of sorting purity. Target cells exhibiting inhibition of NF-κB phenotype were enriched from 67% in the pre-sorting sample to 91% in the sorted sample. **(C)** Evaluation of the reproducibility in training classifiers. The average of the AUC scores was 0.98 ± 0.02 ($n$ = 7). Mean ± SD of the SVM scores are presented and the $Z$'-factor value was obtained as 0.83 ($n$ = 7).

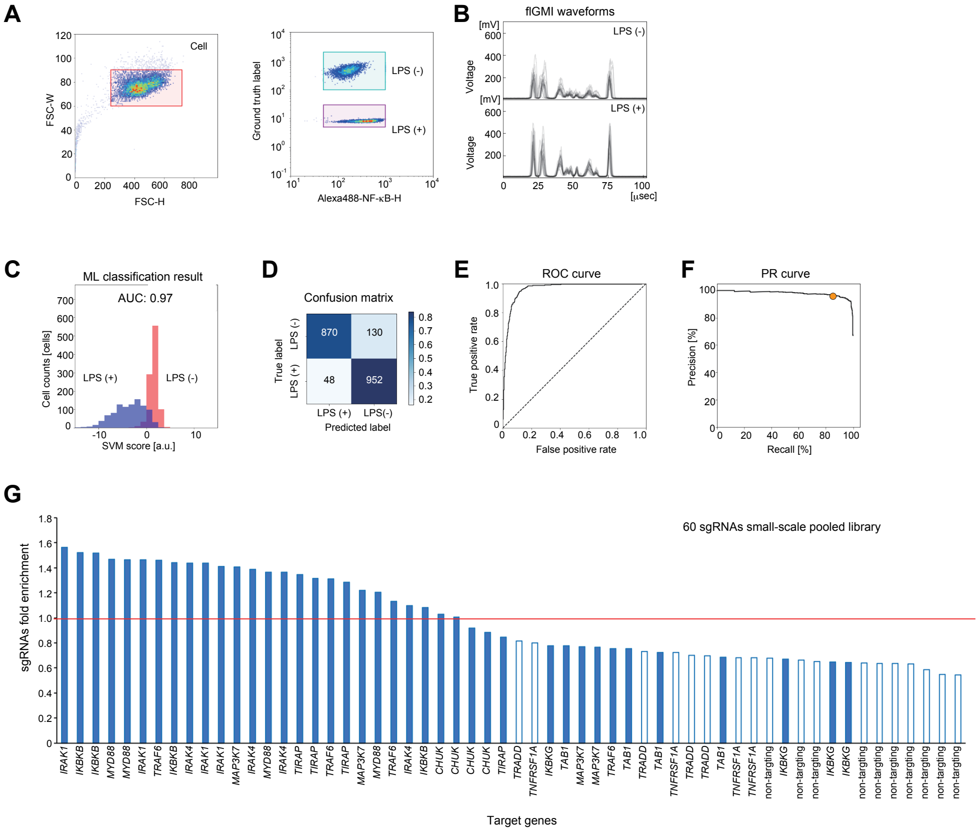


**Fig. S5.**

Data obtained in the pooled CRISPR screening of a small-scale (60 sgRNAs) library by using MViCS in the fluorescence mode (N2: a sample different from N1 but analyzed with the same methods). **(A)** Gating was first performed by the cell gate (left) and then by the fluorescent intensity of the NF-κB-staining and a ground truth marker (Fixable Far-Red dye) which only labeled LPS unstimulated cells (right). **(B)** Randomly selected GMI waveforms for 20 cells in each condition. **(C)** SVM score histogram and AUC score for the classification of LPS (-) and LPS (+) cells. **(D)** Confusion matrix of the classification result. **(E)** ROC curve. **(F)** PR curve. A yellow circle indicates the threshold of 0 < SVM score. **(G)** Fold enrichment of sgRNAs in the small-scale pooled library.

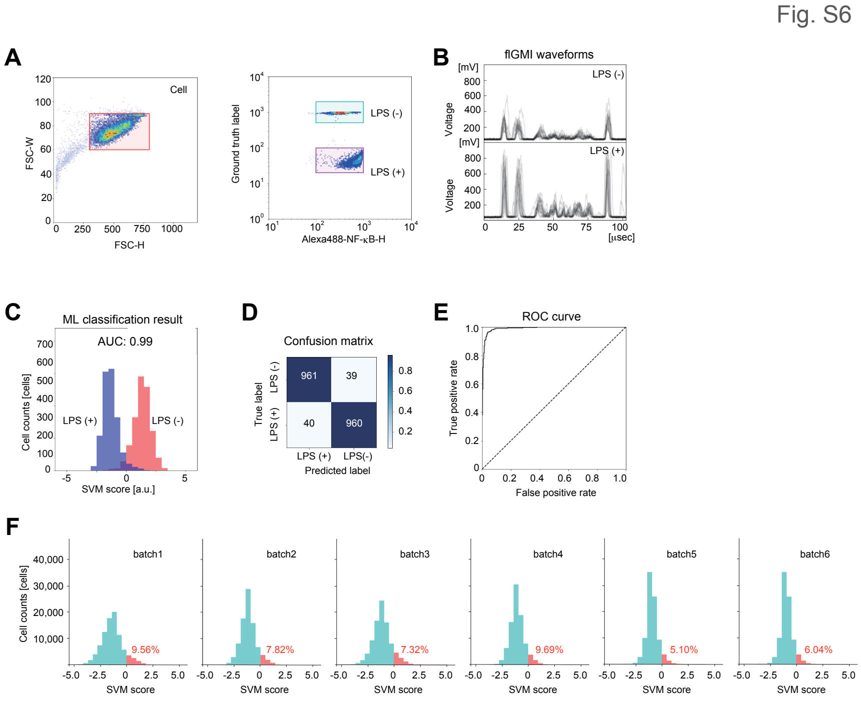


**Fig. S6.**

Data obtained in the pooled CRISPR screening of a kinase library (7,290 sgRNAs) by using MViCS in the fluorescence mode (N1: a sample used in Fig. 3C). **(A)** Gating was first performed by the cell gate (left) and then by fluorescence intensity of the NF-κB-staining and a ground truth marker (Fixable Far-Red dye) which only labeled LPS unstimulated cells (right). **(B)** Randomly selected GMI waveforms of 20 cells in each condition. **(C)** SVM score histogram and AUC score for the classification of LPS (-) and LPS (+) cells. **(D)** Confusion matrix of the classification result. **(E)** ROC curve. **(F)** Percentage of cells predicted as positive (0 < SVM score) during sorting. Batch 1: 1~100,000 cells, batch 2: 100,001~200,000 cells, batch 3: 200,001~300,000 cells, batch 4: 300,001~400,000 cells, batch 5: 400,001~500,000 cells, and batch 6: 500,001~600,000 cells.

Fig. S7

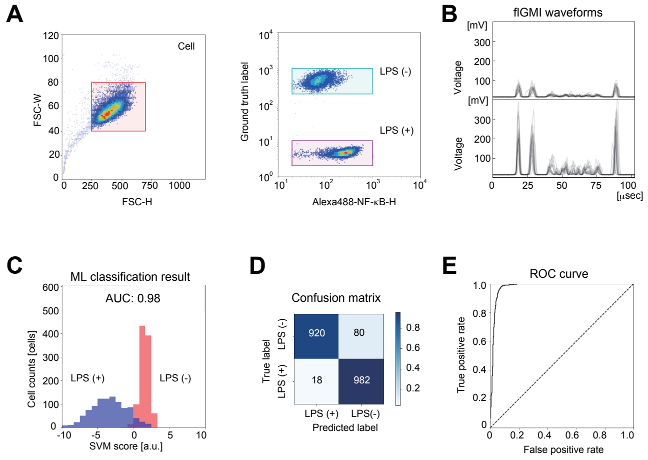


**Fig. S7.**

Data obtained in the pooled CRISPR screening of a kinase library (7,290 sgRNAs) by MViCS in the fluorescence mode sorting (N2: a sample different from N1 but analyzed with the same methods). **(A)** Gating was first performed by the cell gate (left) and then by fluorescence intensity of the NF-κB-staining and a ground truth marker (Fixable Far-Red dye) which only labeled LPS unstimulated cells (right). (**B)** Randomly selected GMI waveforms of 20 cells in each condition. (**C)** SVM score histogram, and AUC score for the classification of LPS (-) and LPS (+) cells. (**D)** Confusion matrix of the classification result. (**E)** ROC curve.

Fig. S8

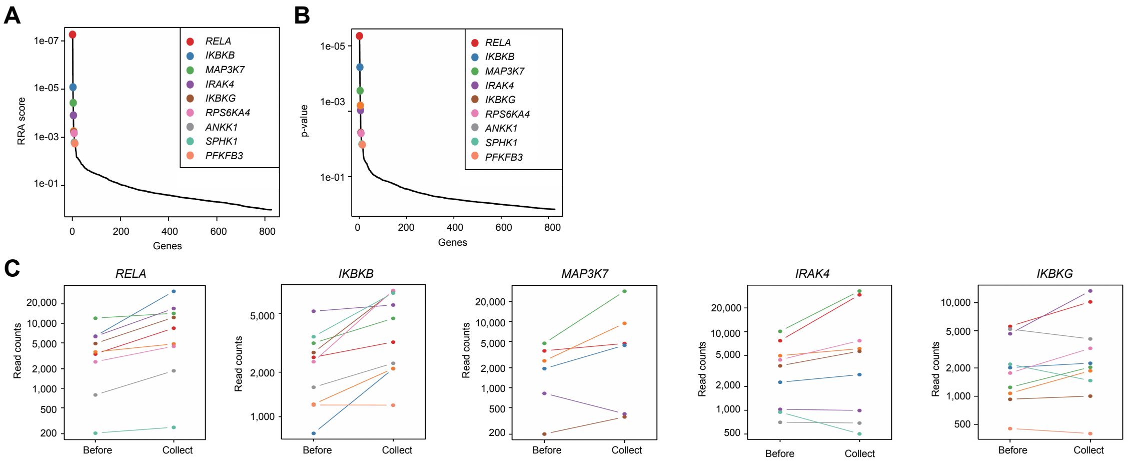


**Fig. S8.**

Results of pooled CRISPR screening by using MViCS in the fluorescence mode (N1: a sample used in Fig. 3C) **(A)** Roust Ranking Aggregation **(**RRA) scores of top 10 hit genes. **(B)** *p*-values of top genes. **(C)** Read counts of sgRNAs for representative hit genes before sorting and their enrichments in the collected sorted cells.

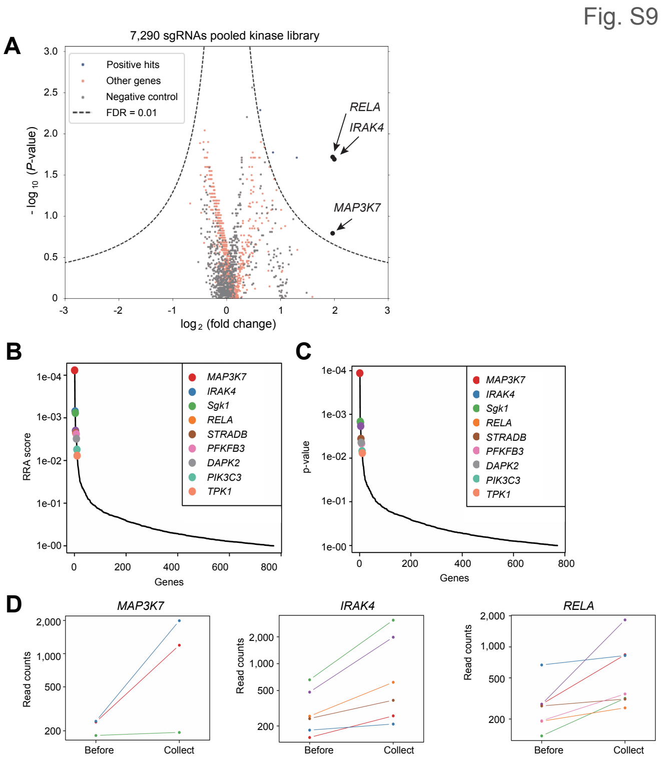


**Fig. S9.**

Results of pooled CRISPR screening using MViCS in the fluorescence mode (N2: a sample different from N1 but analyzed with the same methods) **(A)** Volcano plot visualization of statistical significance (y-axis) and magnitude of the change (x-axis) between before and after the cell sorting, wherein the statistical significance was calculated with Mann-Whitney U test. Dashed lines: cutoff for hit genes (FDR = 0.01). **(B)** RRA scores of the top 10 hit genes. **(C)** *p*-values of top hit genes. **(D)** Read counts of sgRNAs for representative hit genes before sorting and their enrichments in the collected sorted cells.

Fig. S10

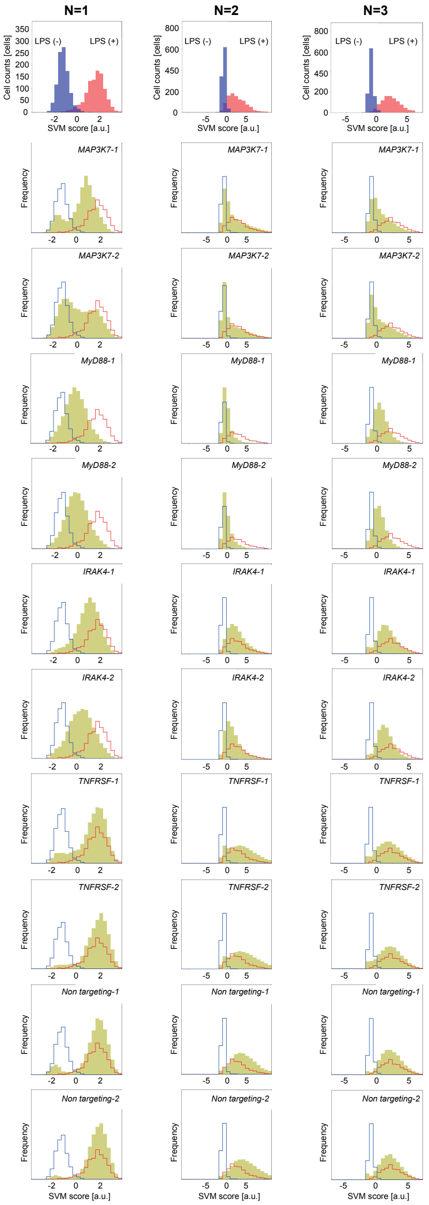


**Fig. S10.**

Data used in Fig. 3E. The three columns of histograms were obtained for three series of independent experiments, respectively. $n$ = 3 biological replicates (N1, N2, and N3). In the top row, the SVM scores were obtained when we developed models for classifying the nuclear translocation using the training samples. From the second top to bottom rows, yellow histograms overlaid on top of the histogram of the first row are the SVM scores obtained when the trained model was applied to cells of each condition: *MAP3K7-1*, *MAP3K-2*, *MYD88-1*, *MYD88-2*, *IRAK4-1*, *IRAK4-2*, *TNFRSF-1*, *TNFRSF-2*, and non-targeting sgRNAs CRISPR knockout cells.

Fig. S11

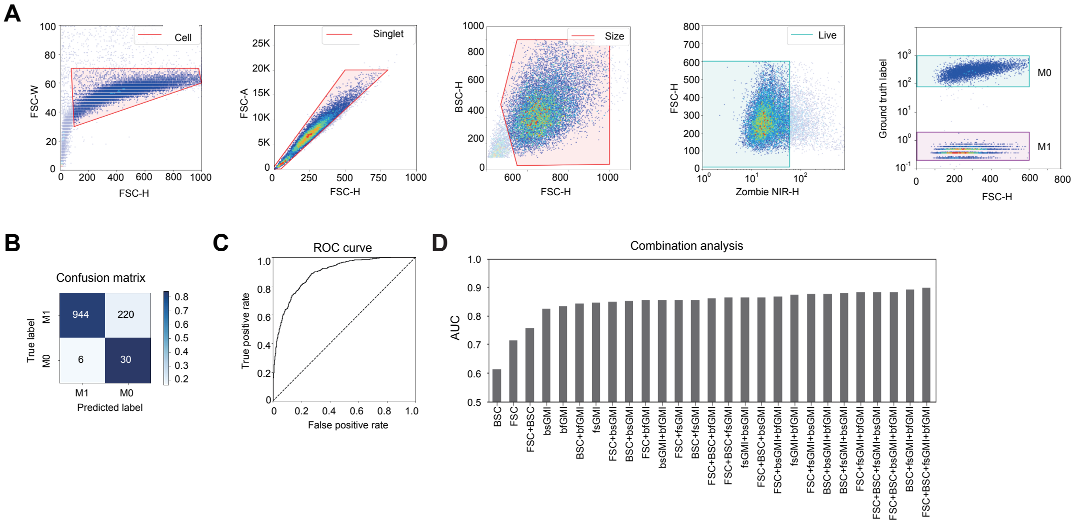


**Fig. S11.**

The development and performance of a machine learning-based model which classifies the high-content phenotypes of M0 and M1 macrophage based on label-free GMI waveforms. **(A)** Gating scheme: cell gate, singlet gate, size gate, live gate, and FSC/Ground truth marker plot of the training sample. M0 macrophages were labeled with a ground truth marker (CellTracker Green dye). **(B)** Confusion matrix for the classification of M0 and M1 polarized macrophages. **(C)** ROC curve. (**D**) Label-free GMI waveforms combination analysis for classification of M0 and M1 macrophages.

Fig. S12

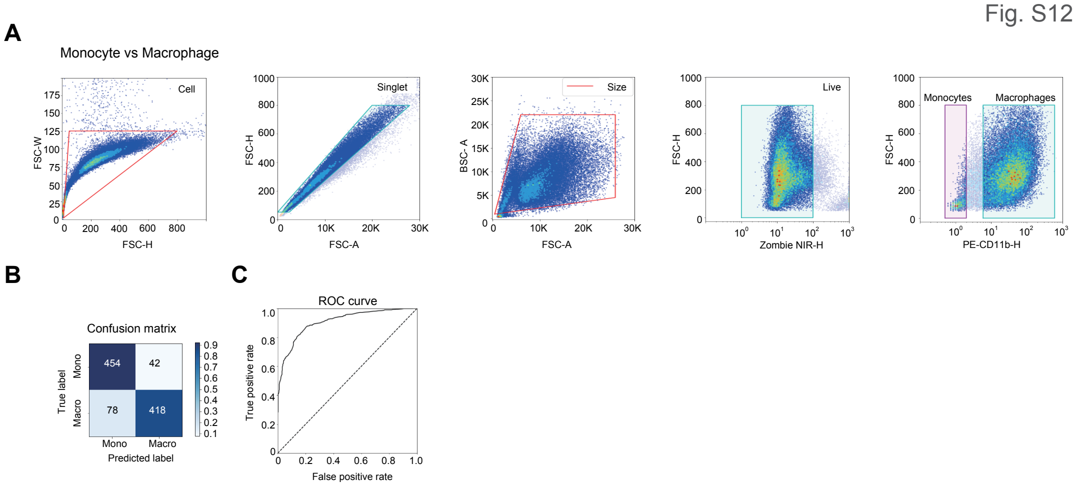


**Fig. S12.**

The development and performance of a machine learning-based model which classifies the high-content phenotypes of THP-1 monocytes and THP-1-derived macrophages based on label-free GMI waveforms. **(A)** Gating scheme; cell gate, singlet gate, size gate, live gate, and CD11b/FSC gate. THP-1-derived CD11b positive cells were identified as positive and CD11b negative cells were identified as negative for a training data set. **(B)** Confusion matrix of the classification result. **(C)** ROC curve.

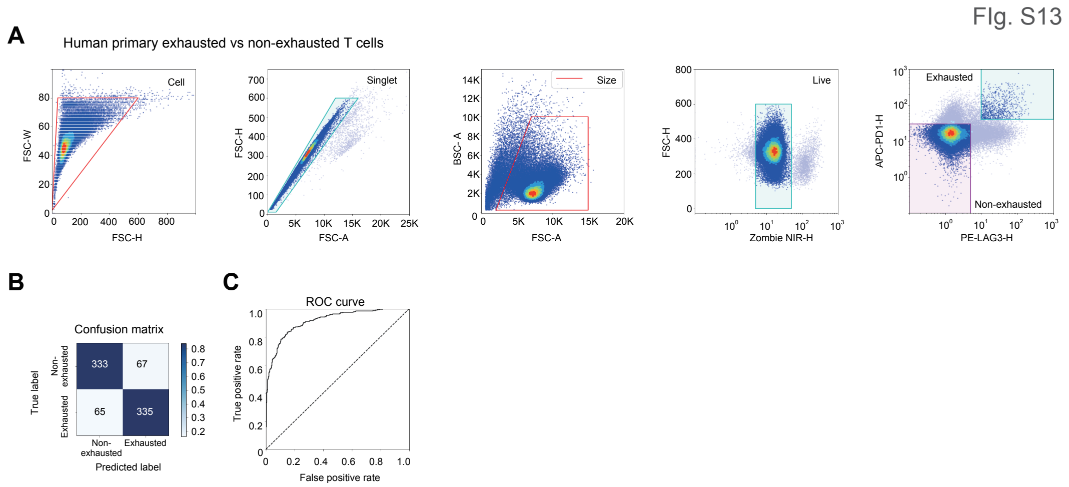


**Fig. S13.**

The development and performance of a machine learning-based model which classifies the high-content phenotypes of exhausted (LAG3/PD-1 double positive) and non-exhausted (LAG3/PD-1 double negative) human primary T cells based on label-free GMI waveforms. **(A)** Gating scheme; cell gate, singlet gate, size gate, live gate, and LAG3/PD-1 gate. LAG3/PD-1 double positive cells were identified as positive and LAG3/PD-1 double negative cells were identified as negative for a training data set. **(B)** Confusion matrix of the classification result. **(C)** ROC curve.

Fig. S14

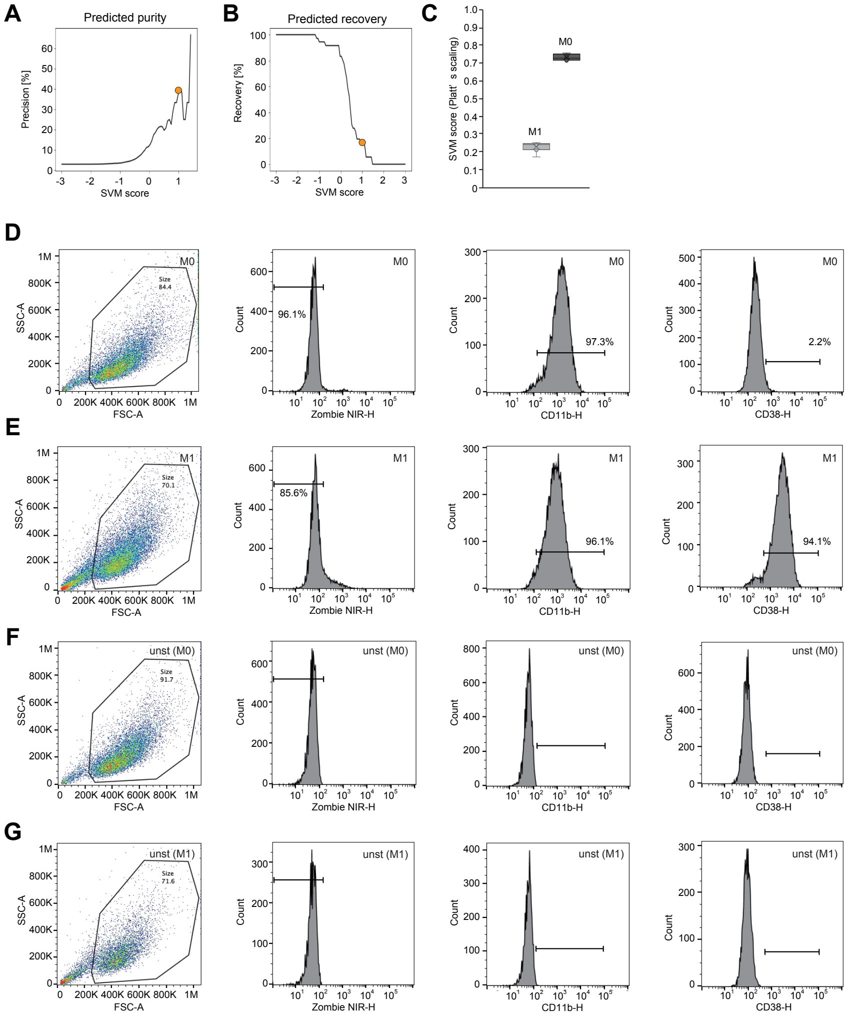


**Fig. S14.**

Data obtained in the pooled CRISPR screening using MViCS in the label-free mode. (N1: a sample used in Figs. 5A). (**A-B)** Assuming the percentage of target cells in the sample was 3 %, (**A)** purity in the sorted sample was expected to be 37.5 %. A yellow circle indicates the threshold of 1 < SVM score. (**B)** Predicted recovery in the sorted sample was 16.7 %. A yellow circle indicates the threshold of 1 < SVM score. (**C)** Evaluation of the reproducibility in training models. The average of the AUC score was 0.85 ± 0.07 (*n* = 7). Mean ± SD of the SVM scores are presented and the Z'-factor value was obtained as 0.75 (*n* = 7). (**D-G)** Validation of the training sample using CD markers. **D,** CD marker-stained M0 cells, **E,** CD marker-stained M1 cells, **F,** unstained M0 cells, **G,** unstained M1 cells.

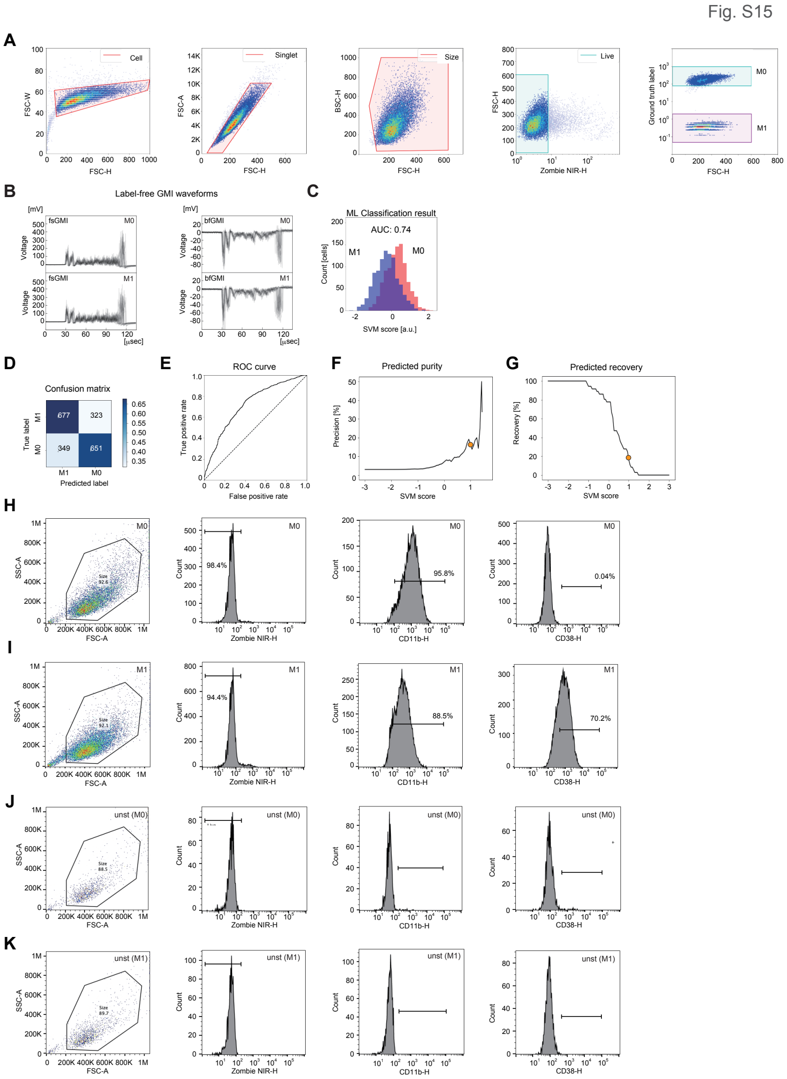


**Fig. S15.**

Data obtained pooled CRISPR screening using MViCS in the label-free mode (N2: a sample different from N1 but analyzed with the same). (**A**) Gating scheme: cell gate, singlet gate, size gate, live gate, and FSC/Ground truth marker plot of the training sample. M0 macrophages were labeled with a ground truth marker (CellTracker Green dye). (**B**) Randomly selected fsGMI and bfGMI waveforms of 20 cells in each population. (**C**) SVM score histogram for the classification of M0 and M1 macrophages. (**D**) Confusion matrix for the classification of M0 and M1 macrophages. (**E**) ROC curve. (**F-G**) Assuming the percentage of target cells in the pre-sorting sample was 3 %, (**F**) purity in the sorted sample was expected to be 16.7 %. A yellow circle indicates the threshold of 1 < SVM score. (**G**) Predicted recovery in the sorted sample was 16.7 %. A yellow circle indicates the threshold of 1 < SVM score. (**H-K**) Validation of the training sample using CD markers. **H,** CD marker-stained M0 cells, **I,** CD marker-stained M1 cells, **J** unstained M0 cells, **K,** unstained M1 cells.

Fig. S16

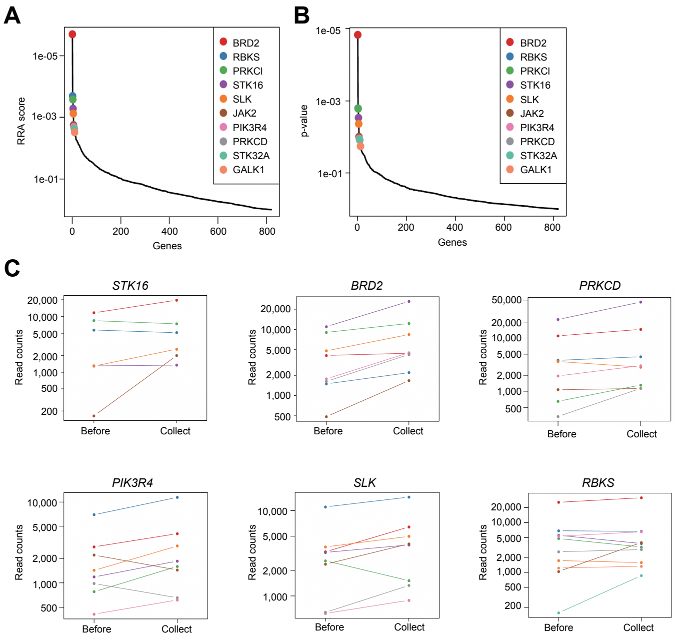


**Fig. S16.**

Results of pooled CRISPR screening using MViCS in the label-free mode (N1: a sample used in Fig. 5A) **(A)** RRA scores of the top 10 hit genes. **(B)** *p*-values of the top 10 hit genes. **(C)** Read counts of sgRNAs for the representative hit genes before sorting and their enrichments in the collected sorted cells.

Fig. S17

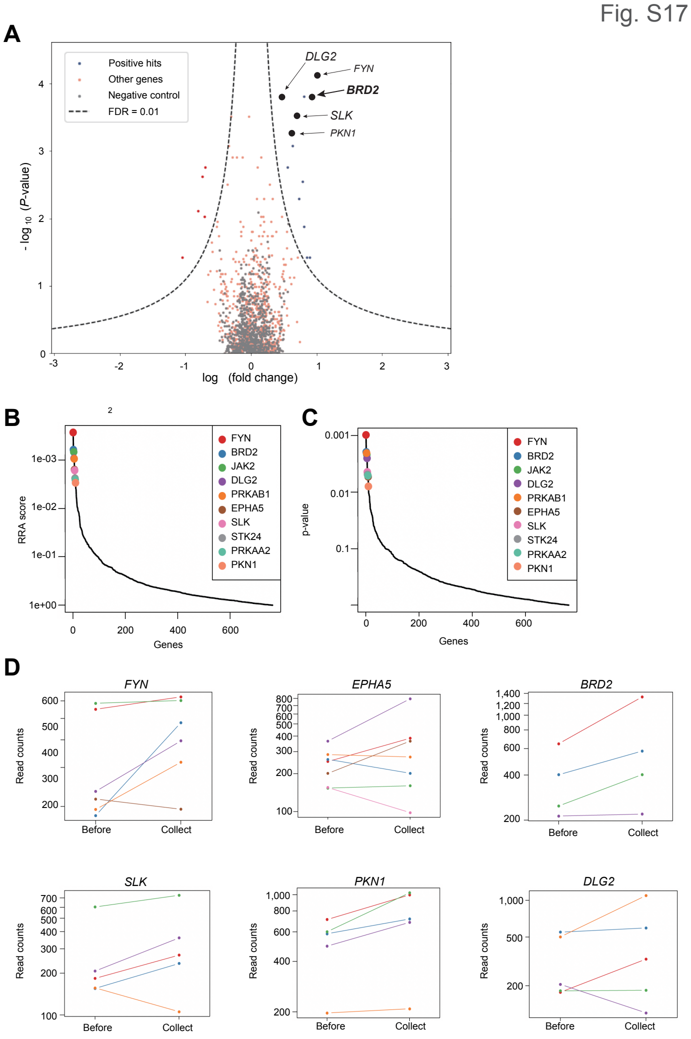


**Fig. S17.**

Results of pooled CRISPR screening using MViCS in the label-free mode (N2: a sample different from N1 but analyzed with the same) **(A)** Volcano plot visualization of statistical significance (y-axis) and magnitude of the change (x-axis) between before and after the cell sorting, wherein the statistical significance was calculated with Mann-Whitney U test. Dashed lines: cutoff for hit genes (FDR = 0.01). **(B),** RRA scores of the top 10 hit genes. **(C)** *p*-values of top 10 hit genes. **(D)** Read counts of sgRNAs for the representative hit genes before sorting and their enrichments in the collected sorted cells.

Fig. S18

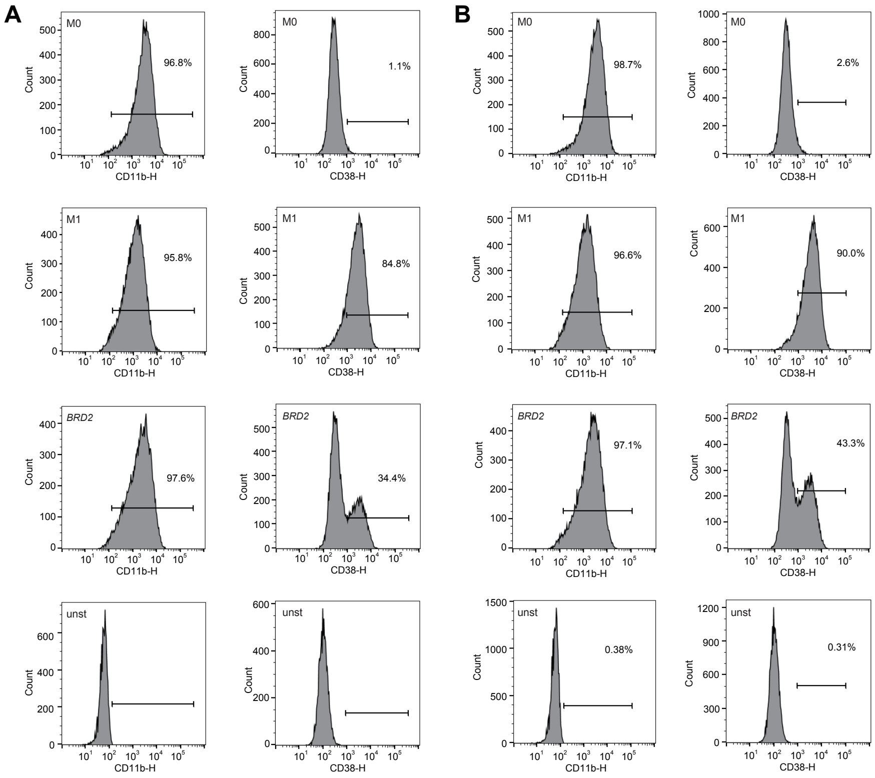


**Fig. S18.**

Macrophage surface marker expression of control M0, control M1, and *BRD2* CRISPR KO M1 cells. Samples were different from the one used in Fig. 5B but analyzed with the same methods (**A**: N2, **B:** N3). CD11b was used as a pan-macrophage marker, and CD38 was used as an M1-specific marker.

Table.S1

| gRNA_name | Target_gene | gRNA_seqeunce |
|---|---|---|
| TNFRSF1A-1 | TNFRSF1A | CAGCAGCAGGTCAGGCACGG |
| TNFRSF1A-2 | TNFRSF1A | TGGCAGCAGCAGGTCAGGCA |
| TNFRSF1A-3 | TNFRSF1A | GCAGGTCAGGCACGGTGGAG |
| TNFRSF1A-4 | TNFRSF1A | GTCTCACCAGTGGCAGCAGC |
| TRADD-1 | TRADD | CAGGACCACCTTGTCCAGCG |
| TRADD-2 | TRADD | ACACTGCCACCTTCTGCTGG |
| TRADD-3 | TRADD | AATGGGCACGAAGAGTGGGT |
| TRADD-4 | TRADD | GGAGTCCTCGCTGGACAAGG |
| MYD88-1 | MYD88 | CGCATGTTGAGAGCAGCCAG |
| MYD88-2 | MYD88 | CGTTCAAGAACAGAGACAGG |
| MYD88-3 | MYD88 | GCGGTCCTGGAGCCTCAGCG |
| MYD88-4 | MYD88 | CCGGCAACTGGAGACACAAG |
| IRAK1-1 | IRAK1 | AAGCGGCACATGACCCAGGG |
| IRAK1-2 | IRAK1 | TAGAAGCGGCACATGACCCA |
| IRAK1-3 | IRAK1 | CACCTCGTACAAGAAGTGCT |
| IRAK1-4 | IRAK1 | CGCCCAGCACTTCTTGTACG |
| IRAK4-1 | IRAK4 | TTGTATCTATCATCACCAGA |
| IRAK4-2 | IRAK4 | GATTTTATTGATCCTCAAGA |
| IRAK4-3 | IRAK4 | TCCTAATTAGTCCAACATTG |
| IRAK4-4 | IRAK4 | GTAGCTATTAAAAAACCATC |
| TRAF6-1 | TRAF6 | GTAACAAAAGATGATAGTGT |
| TRAF6-1 | TRAF6 | GTCTGAAAGTGACTGCTGTG |
| TRAF6-3 | TRAF6 | TGGGTGGAACTGCCAGCACG |
| TRAF6-4 | TRAF6 | GTAATGCCATCAAGCAGATG |
| TIRAP-1 | TIRAP | TCTCGGCCTAAGAAGCCTCT |
| TIRAP-2 | TIRAP | GCCGAGAGCCAGGAGCTGGG |
| TIRAP-3 | TIRAP | TAAGAAGCCTCTAGGCAAGA |
| TIRAP-4 | TIRAP | AGAGCCAGGAGCTGGGAGGG |
| MAP3K7-1 | MAP3K7 | CGACTACAAGGAGATCGAGG |
| MAP3K7-2 | MAP3K7 | GATCGACTACAAGGAGATCG |
| MAP3K7-3 | MAP3K7 | AGGGGCTTCGATCATCTCAC |
| MAP3K7-4 | MAP3K7 | CTCACCGGCCGAAGACGAGG |
| CHUK-1 | CHUK | ACAGACGTTCCCGAAGCCGC |
| CHUK-2 | CHUK | TGTACCAGCATCGGGTGAGG |
| CHUK-3 | CHUK | CGTCTGTCTGTACCAGCATC |
| CHUK-4 | CHUK | ACGTCTGTCTGTACCAGCAT |
| IKBKB-1 | IKBKB | CCTGACAACGCAGACATGTG |
| IKBKB-2 | IKBKB | AAAGAGCGCCTTGGGACAGG |
| IKBKB-3 | IKBKB | GATTTGGAAATGTCATCCGA |
| IKBKB-4 | IKBKB | CCCACATGTCTGCGTTGTCA |
| IKBKG-1 | IKBKG | GAAGCTCTCCAGCATCATCG |
| IKBKG-2 | IKBKG | ACTGGGATTGATAGTCTGCG |
| IKBKG-3 | IKBKG | TTTCCCCACCTGGATCACAA |
| IKBKG-4 | IKBKG | AACTAAGGCCCAGAGAAGTG |
| TAB1-1 | TAB1 | AGGCTGAGCCAACCCCAGAG |
| TAB1-2 | TAB1 | ACCCCAGAGAGGTGGCAGAG |
| TAB1-3 | TAB1 | CAGGGGTTCCTCCAAGATGG |
| TAB1-4 | TAB1 | AGCTACTCTGCTGATGGCAA |
| non targeting | Scramble | GCACTACCAGAGCTAACTCA |
| non targeting | EGFP | GGCCACAAGTTCAGCGTGTC |
| non targeting | EGFP | GGGCGAGGAGCTGTTCACCG |
| non targeting | EGFP | GCCGTCCAGCTCGACCAGGAT |
| non targeting | mCherry | GGCCACGAGTTCGAGATCGA |
| non targeting | mCherry | GGCGAGGGCCGCCCCTACGA |
| non targeting | mCherry | GCGGTGTGGGTGCCCTCGTA |
| non targeting | AAVS1 | GAACCTGAGCTGCTCTGACG |
| non targeting | AAVS1 | GTTGGCAGGGGGTGGGAGGGA |
| non targeting | AAVS1 | GGGAGGGAGAGCTTGGCAGG |
| non targeting | AAVS1 | GCCTCTAAGGTTTGCTTACGA |
| non targeting | AAVS1 | GCTCCCTCCCAGGATCCTCTC |
| BRD2-1 | BRD2 | TTGGCCTTTCTATAAACCAG |
| BRD2-2 | BRD2 | GTGGGGAATGTTGAGGACAG |
| BRD2-3 | BRD2 | TAAGAACAGCCACAAGAAGG |
| BRD2-4 | BRD2 | AGGTAGTGATGAAGGCTCTG |

**Table S1.**

A list of the small-scale sgRNA sequences.

Table. S2

| Gene | Epsilon | p-value | Product |
|---|---|---|---|
| STK16 | -1.874746667 | 5.42593597395551E-05 | -7.996779404 |
| BRD2 | -1.565743333 | 1.35648399348888E-05 | -7.621389274 |
| LRRK1 | -1.50134 | 5.42593597395551E-05 | -6.404003807 |
| DLG2 | -1.31701 | 2.71296798697775E-05 | -6.014199036 |
| PRKCD | -1.366076667 | 9.49538795442214E-05 | -5.495026024 |
| TESK2 | -1.326283333 | 9.49538795442214E-05 | -5.334957847 |
| PIM2 | -1.410243333 | 0.000217037 | -5.166377578 |
| MAPKAPK2 | -1.407343333 | 0.000217037 | -5.155753529 |
| CDK8 | -1.24284 | 0.000149213 | -4.755345262 |
| PIK3CD | -1.178463333 | 9.49538795442214E-05 | -4.740353776 |
| TYRO3 | -1.276653333 | 0.000217037 | -4.676975243 |
| LIMK1 | -1.21189 | 0.000149213 | -4.6369246 |
| IRAK3 | -1.56764 | 0.001383614 | -4.481859488 |
| MATK | -1.073223333 | 9.49538795442214E-05 | -4.317027214 |
| PIK3CB | -1.142426667 | 0.000217037 | -4.185240501 |
| BMP2K | -1.441616667 | 0.001383614 | -4.121560649 |
| RPS6KC1 | -1.098983333 | 0.000217037 | -4.026087355 |
| Cdk11b | -1.42366 | 0.001994031 | -3.844263529 |
| STRADA | -1.15781 | 0.000556158 | -3.76844169 |
| PIK3R4 | -1.000023333 | 0.000217037 | -3.663550825 |
| CAMK2B | -1.167766667 | 0.000908844 | -3.551774606 |
| ERN1 | -1.214706667 | 0.001383614 | -3.472828327 |
| SLK | -1.016743333 | 0.00042051 | -3.43275287 |
| VRK3 | -1.099863333 | 0.001668475 | -3.05506862 |
| RBKS | -1.649813333 | 0.015423223 | -2.989172815 |
| EPHA5 | -1.10022 | 0.001994031 | -2.97088885 |
| TAOK2 | -1.014843333 | 0.001383614 | -2.901422024 |
| PRKCI | -1.063916667 | 0.001994031 | -2.872860121 |
| STK24 | -1.12226 | 0.002767227 | -2.870690757 |
| TAF1 | -0.944376667 | 0.001125882 | -2.784501433 |
| KIAA1804 | -1.16744 | 0.004856213 | -2.701108613 |
| ILK | -1.03604 | 0.002767227 | -2.650143863 |
| MYLK2 | -0.951523333 | 0.001994031 | -2.569368 |
| BLK | -1.083663333 | 0.00425936 | -2.568992635 |
| PPAN | -1.127143333 | 0.005507325 | -2.546283622 |
| ABL1 | -1.012163333 | 0.003214867 | -2.523158188 |
| DYRK2 | -0.874176667 | 0.001383614 | -2.499258113 |
| POMK | -1.28757 | 0.011733587 | -2.485743009 |
| EEF2K | -0.728883333 | 0.00042051 | -2.460873135 |
| ALPK3 | -0.937633333 | 0.002767227 | -2.398424022 |
| PRKAR1A | -0.916523333 | 0.003214867 | -2.284743259 |
| YAP1 | -0.888693333 | 0.002767227 | -2.273237697 |
| HIPK3 | -0.972633333 | 0.004856213 | -2.25038398 |
| TNK2 | -0.80367 | 0.001668475 | -2.232338258 |
| LAMB1 | -0.990686667 | 0.006212697 | -2.186167931 |
| THNSL1 | -0.869743333 | 0.003214867 | -2.168128345 |
| MAPK12 | -0.884463333 | 0.003716766 | -2.149099755 |
| CDC7 | -0.705086667 | 0.000908844 | -2.144528517 |
| MAP4K3 | -0.93567 | 0.005507325 | -2.113734009 |
| MAP3K3 | -0.940456667 | 0.006212697 | -2.075324393 |

**Table S2.**

Top 50 hit genes obtained as a result of the pooled CRISPR screening MViCS in the label-free mode (N1: a sample used in Fig. 5A).

Table. S3

N2

| Genes | epsilon | pvalue | product |
|---|---|---|---|
| FYN | -1.01323 | 7.62340384981895E-05 | -4.172330243 |
| EPHA5 | -0.924293333 | 0.000152468 | -3.527862276 |
| BRD2 | -0.8103 | 0.000152468 | -3.092770119 |
| SLK | -0.709763333 | 0.000304936 | -2.495379597 |
| PKN1 | -0.637376667 | 0.000533638 | -2.085976419 |
| PRKAB1 | -0.784698333 | 0.002795248 | -2.003789689 |
| MAP4K2 | -0.628636667 | 0.000838574 | -1.933974546 |
| DLG2 | -0.47333 | 0.000152468 | -1.80661592 |
| AK7 | -0.730786667 | 0.005107681 | -1.674799541 |
| SGK223 | -0.555386667 | 0.001753383 | -1.530714097 |
| IRAK4 | -0.801953333 | 0.013264723 | -1.505508456 |
| DBF4 | -0.896183333 | 0.037735849 | -1.275489231 |
| PIK3R4 | -0.850783333 | 0.037735849 | -1.210873869 |

N3

| Genes | epsilon | pvalue | product |
|---|---|---|---|
| IKBKB | -0.772620667 | 5.2578999947421E-05 | -3.306188837 |
| OXSR1 | -0.669340667 | 0.000814974 | -2.067496921 |
| ANKK1 | -0.554800667 | 0.000420632 | -1.87306125 |
| PIK3R2 | -0.649894 | 0.001393343 | -1.856059443 |
| MARK2 | -0.566397333 | 0.000814974 | -1.74951979 |
| TRIB1 | -0.546360667 | 0.000814974 | -1.687629413 |
| Cdk16 | -0.582167333 | 0.001393343 | -1.662636024 |
| CDKN1A | -0.557377333 | 0.001077869 | -1.653980348 |
| PIK3R3 | -0.539104 | 0.001393343 | -1.539649651 |
| NEK11 | -0.598587333 | 0.004574373 | -1.400495884 |
| MAP4K2 | -0.507174 | 0.002681529 | -1.304257535 |
| STK40 | -0.672200667 | 0.024975025 | -1.077197581 |
| BTK | -0.300574 | 0.000289184 | -1.063678782 |
| Tab1 | -0.490014 | 0.009411641 | -0.992932346 |
| PIP5K1A | -0.451250667 | 0.009411641 | -0.914384861 |
| MAPK7 | -0.449454 | 0.012040591 | -0.862660522 |
| MAPK11 | -0.425537333 | 0.012040591 | -0.816756015 |
| CDK7 | -0.475177333 | 0.020637257 | -0.800839177 |
| BRD2 | -0.003014 | 0.339712919 | -0.001413228 |
| CDK10 | -0.079317333 | 0.961038961 | -0.001368938 |

**Table S3.**

List of hit genes obtained as a result of the pooled CRISPR screening using MViCS in the label-free mode. Samples are different from the one used in Fig. 5A but analyzed with the same methods (N2: top, N3: bottom).

Table. S4

1st PCR

| | | |
|---|---|---|
| small scale | Forward | TAACTTACGGAGTCGCTCTACGtcttgtggaaaggacgaaacacc |
| | Reverse | GGATGGGATTCTTTAGGTCCTGttgtctcaagatctagttacgc |
| large scale | Forward | TAACTTACGGAGTCGCTCTACGcgtaacttgaaagtatttc |
| | Reverse | GGATGGGATTCTTTAGGTCCTGcaagttgataacggactag |

2nd PCR

| | | |
|---|---|---|
| index1 | Forward | AATGATACGGCGACCACCGAGATCTACACTCTTTCCCTACACGACGCTCTTCCGATCT NNNNN CAAGTGTTC TAACTTACGGAGTCGCTCTACG |
| | Reverse | CAAGCAGAAGACGGCATACGAGATCGGTCTCGGCATTCCTGCTGAACCGCTCTTCCGATCT NNNNN CAAGTGTTC GGATGGGATTCTTTAGGTCCTG |
| index2 | Forward | AATGATACGGCGACCACCGAGATCTACACTCTTTCCCTACACGACGCTCTTCCGATCT NNNNN AGGACATTC TAACTTACGGAGTCGCTCTACG |
| | Reverse | CAAGCAGAAGACGGCATACGAGATCGGTCTCGGCATTCCTGCTGAACCGCTCTTCCGATCT NNNNN AGGACATTC GGATGGGATTCTTTAGGTCCTG |
| index3 | Forward | AATGATACGGCGACCACCGAGATCTACACTCTTTCCCTACACGACGCTCTTCCGATCT NNNNN CACTAATGG TAACTTACGGAGTCGCTCTACG |
| | Reverse | CAAGCAGAAGACGGCATACGAGATCGGTCTCGGCATTCCTGCTGAACCGCTCTTCCGATCT NNNNN CACTAATGG GGATGGGATTCTTTAGGTCCTG |
| index4 | Forward | AATGATACGGCGACCACCGAGATCTACACTCTTTCCCTACACGACGCTCTTCCGATCT NNNNN AGCCTGATG TAACTTACGGAGTCGCTCTACG |
| | Reverse | CAAGCAGAAGACGGCATACGAGATCGGTCTCGGCATTCCTGCTGAACCGCTCTTCCGATCT NNNNN AGCCTGATG GGATGGGATTCTTTAGGTCCTG |

**Table S4.**

PCR primers for amplicon sequencing.
